# Physiological and genomic analysis of “*Candidatus* Nitrosocosmicus agrestis”, an ammonia tolerant ammonia-oxidizing archaeon from vegetable soil

**DOI:** 10.1101/2019.12.11.872556

**Authors:** Liangting Liu, Mengfan Liu, Yiming Jiang, Weitie Lin, Jianfei Luo

## Abstract

The presences of ammonia tolerant ammonia-oxidizing archaea (AOA) in environments are always underestimated and their adaption to complex habitats has also rarely been reported. Here we present the physiological and genomic characteristics of an ammonia tolerant soil AOA strain *Candidatus* Nitrosocosmicus agrestis. This strain was able to form aggregates and adhere on the surface of hydrophobic matrix. Ammonia-oxidizing activities were still observed at 200 mM NH_4_^+^ (> 1500 μM of free ammonia) and 50 mM NO_2_^-^. Urea could be used as sole energy source but exogenous organics had no significant effect on the ammonia oxidation. Besides the genes involving in ammonia oxidation, carbon fixation and urea hydrolysis, the genome also encodes a full set of genes (GTs, GHs, CEs, MOP, LPSE, etc) that responsible for polysaccharide metabolism and secretion, suggesting the potential production of extracellular polymeric substances (EPS). Moreover, a pathway connecting urea cycle, polyamines synthesis and excretion was identified in the genome, which indicates the NH_4_^+^ in cytoplasm could potentially be converted into polyamines and excreted out of cell, and then contributes to the high ammonia tolerance. Genes encoding the cytoplasmic carbonic anhydrase and putative polyamine exporter are unique in *Ca*. Nitrosocosmicus agrestis or the genus *Ca*. Nitrosocosmicus, suggesting the prevalence of ammonia tolerance in this clade. The proposed mechanism of ammonia tolerance via polyamines synthesis and export was verified by using transcriptional gene regulation and polyamines determination.

**IMPORTANCE:** AOA are ubiquitous in different environments and play a major role in nitrification. Though AOA have higher affinities for ammonia, their maximum specific cell activity and ammonia tolerance are usually much lower than AOB, resulting in low contribution to the global ammonia oxidation and N_2_O production. However, in some agricultural soils, the AOA activity would not be suppressed by the fertilization with high concentration of ammonium nitrogen, suggesting the presence of some ammonia tolerant species. This study provides some physiological and genomic characteristics for an ammonia tolerant soil AOA strain *Ca*. Nitrosocosmicus agrestis and proposes some mechanisms of this AOA adapting to a variety of environments and tolerating to high ammonia. Ammonia tolerance of AOA was always underestimated in many previous studies, physiological and genomic analyses of this AOA clade are benefit to uncover the role of AOA playing in global environmental patterns.

## INTRODUCTION

Ammonia-oxidizing archaea (AOA) of the phylum *Thaumarchaeota* play a major role in nitrification that mediating the conversion of ammonia to nitrate via nitrite (Stahl and de la Torre, 2012; Stein and Klotz, 2016; Kuypers *et al*., 2018; Stein, 2019). AOA are ubiquitous and able to thrive in different environments. It was estimated that the AOA represents up to 1-5% of all prokaryotes in soils and 20–40% of all marine bacterioplankton in marine system (Stahl and de la Torre, 2012). AOA within the phylum *Thaumarchaeota* is composed of four orders (Kerou *et al*., 2016): (i) *Ca*. Nitrosopumilales: mainly (*Ca*. Nitrosopumilus, *Ca*. Nitrosopelagicus) derived from the oligotrophic ocean (Könneke *et al*., 2005; Hatzenpichler, 2012; Qin *et al*., 2014; Santoro *et al*., 2015; Bayer *et al*., 2016) and some of them (*Ca*. Nitrosoarchaeum, *Ca*. Nitrosotenuis) come from terrestrial habitats (Jung *et al*., 2011, 2014; Li *et al*., 2016; Sauder *et al*., 2018). (ii) *Ca*. Nitrosotaleales: represented by the genus *Ca*. Nitrosotalea that mainly distributed in acidic soil (Lehtovirta-Morley *et al*., 2011). (iii) *Ca*. Nitrosocaldales: members of thermophilic AOA belonging to genus *Ca*. Nitrosocaldus (Hatzenpichler *et al*., 2008; Abby *et al*., 2018; Daebeler *et al*., 2018). (iv) *Nitrososphaerales: Nitrososphaera, Ca*. Nitrosocosmicus and many uncultivated AOA that inhabit in soils (Tourna *et al*., 2011; Zhalnina *et al*., 2014; Lehtovirta-Morley *et al*., 2016), sediments (Jung *et al*., 2016), terrestrial hot springs (Hatzenpichler *et al*., 2008) and wastewater treatment plants (Sauder *et al*., 2017). Up to now, four strains belonging to *Nitrososphaera* and five strains belonging to *Ca*. Nitrosocosmicus have been obtained (Alves *et al*., 2019; Liu *et al*., 2019). As a sister cluster of *Nitrososphaera*, the *Ca*. Nitrosocosmicus owns many different properties from *Nitrososphaera* and the other AOA clusters, especially the ability to tolerate high concentration of ammonia (Jung *et al*., 2016; Lehtovirta-Morley *et al*., 2016; Sauder *et al*., 2017). Nitrification is a central process in the global nitrogen cycle. Owing to the ammonia affinities of AOA are dozens or hundreds of times higher than AOB (Könneke *et al*., 2005; Martens-Habbena *et al*., 2009; Kits *et al*., 2017), members of AOA have usually been observed to have higher abundance than AOB in the habitats containing low ammonium, such as the oligotrophic oceans (Berg *et al*., 2015; Trimmer *et al*., 2016). However, in agricultural soils, the overdose fertilization always results in high concentration of ammonia. Though AOA have higher affinities for ammonia, their maximum specific cell activity (*V_max_*) and ammonia tolerance are much lower than AOB (Prosser and Nicol, 2012; Kits *et al*., 2017; Liu et al., 2019). High ammonium in agricultural soils usually contributed to the activation of AOB and the limitation of AOA, even though the AOA groups dominated in numbers (Sterngren *et al*., 2015; Egan *et al*., 2018). Aerobic ammonia oxidation that performed by AOA and AOB is usually regarded as the main source of nitrous oxide (N_2_O) and contributes to the global N_2_O emissions from agriculture (Reay *et al*., 2012; Oertel *et al*., 2016; Lam *et al*., 2017). In general, the AOA species were reported to have low ammonia tolerance (only up to 1-20 mM) (Table 1) while AOB species were able to tolerate 7-50 mM (Prosser and Nicol, 2012) even up to 400 mM (Hunik *et al*., 1992). Then, AOB groups were usually regarded as the main contributor who was responsible for the N_2_O emission in agricultural fields after fertilizing with ammonium nitrogen (Wang *et al*., 2016; Hink *et al*., 2017; Meinhardt *et al*., 2018). However, the AOA groups were still observed to have a small contribution (10-20%) to the N_2_O emission (Wang *et al*., 2016; Hink *et al*., 2017; Meinhardt *et al*., 2018). Moreover, in some agricultural soils, the ammonia oxidation and N_2_O production of AOA would not be suppressed by the fertilization with high ammonium (Schauss *et al*., 2009; Stopnišek *et al*., 2010; Lu *et al*., 2015; Duan *et al*., 2019). It seems that some unknown AOA species was able to tolerate high ammonia in these soils. No evidence indicated that the presence of ammonia tolerant AOA until obtaining the enrichments of *Nitrosocosmicus* from soil, sediment and WWTP (Jung *et al*., 2016; Lehtovirta-Morley *et al*., 2016; Sauder *et al*., 2017).

**Table 1.** Ammonia, nitrite and pH tolerances of AOA strains

| AOA Strain | Source | Inhibitory $\text{NH}_4^+$(mM)<sup>a</sup> / $\text{NH}_3$($\mu\text{M}$)<sup>b</sup> | Inhibitory $\text{NO}_2^-$(mM)<sup>c</sup> | pH<sup>d</sup> | Reference |
| --- | --- | --- | --- | --- | --- |
| **Ca. Nitrosopumilales** |  |  |  |  |  |
| *Nitrosopumilus maritimus* SCM1 | Tropical marine aquarium | 2 / 49.49 | N/A | 6.8-8.2 | (Martens-Habbena *et al.*, 2009) |
| *Ca. Nitrosopumilus koreensis* AR1 | Marine sediment | 4 / 399.04 | N/A | N/A | (Park *et al.*, 2010) |
| *Ca. Nitrosoarchaeum koreensis* MY1 | Agricultural soil | 20 / 112.71 | 10 (7.0) | 6.0-8.0 | (Jung *et al.*, 2011) |
| *Ca. Nitrosotenuis chungbukensis* MY2 | Agricultural soil | 20 / 112.71 | 2 (7.0) | 6.5-8.5 | (Jung *et al.*, 2014) |
| *Ca. Nitrosotenuis cloacae* SAT1 | Wastewater treatment plant | 5 / 11.82 | 5 (6.5) | 5.0-7.0 | (Li *et al.*, 2016) |
| *Nitrosopumilus cobalaminigenes* HCA1 | 50 m depth marine water | 10 / 176.06 | N/A | 6.8-8.1 | (Qin *et al.*, 2017) |
| *Nitrosopumilus ureiphilus* PS0 | Marine surface sediment | 20 / 377.35 | N/A | 5.9-8.1 | (Qin *et al.*, 2017) |
| *Nitrosopumilus oxyclinae* HCE1 | 17 m depth marine water | 1 / 17.61 | N/A | 6.4-7.8 | (Qin *et al.*, 2017) |
| *Ca. Nitrosotenuis aquariensis* AQ6f | Freshwater aquarium biofilter | 3 / 607.18 | N/A | N/A | (Sauder *et al.*, 2018) |
| **Ca. Nitrosotaleales** |  |  |  |  |  |
| *Ca. Nitrosotalea devanattera* Nd1 | Acid soil | 50 / 1.11 | 0.04 (4.5) | 4.0-5.5 | (Lehtovirta-Morley *et al.*, 2011) |
| **Nitrososphaerales** |  |  |  |  |  |
| *Ca. Nitrososphaera gargensis* Ga9.2 | Hot spring | 3 / 161.78 | N/A | N/A | (Hatzenpichler *et al.*, 2008) |
| *Nitrososphaera viennensis* EN76 | Garden soil | 20 / 686.11 | 10 (7.5) | 6.0-8.5 | (Tourn *et al.*, 2011; Stieglmeier *et al.*, 2014) |
| *Ca. Nitrososphaera* sp. JG1 | Agricultural soil | 20 / 80.71 | 10 (6.5) | 6.0-8.0 | (Kim *et al.*, 2012) |
| *Ca. Nitrosocosmicus oleophilus* MY3 | Coal tar-contaminated sediment | 50 / 398.02 | N/A | 5.5-8.5 | (Jung *et al.*, 2016) |
| *Ca. Nitrosocosmicus franklandus* C13 | Arable soil | 100 / 4689.43 | 20 (7.5) | 6-8.5 | (Lehtovirta-Morley *et al.*, 2016) |
| *Ca. Nitrosocosmicus exaquare* G61 | Wastewater treatment plant | 20 / 1485.64 | 30 (8.0) | N/A | (Sauder *et al.*, 2017) |
| *Ca. Nitrososphaera* sp. OTU8 | Wastewater treatment plant | 3 / 23.88 | N/A | 7.0-8.0 | (Chen *et al.*, 2017) |
| *Ca. Nitrosocosmicus agrestis* SS | Vegetable soil | 200 / 1592.07 | 50 (7.0) | 6.0-8.0 | This study |

Inhibitory $\text{NH}_4^+$ (mM)<sup>a</sup>: Ammonium chloride concentration that completely or almost inhibited AOA ammonia oxidization. $\text{NH}_3$ ( $\mu\text{M}$ )<sup>b</sup>: Free ammonia concentration, calculated according to total ammonia, pH and temperature, estimating by Emerson *et al.*'s (Emerson *et al.*, 1975) formulas. Inhibitory $\text{NO}_2^-$ (mM)<sup>c</sup>: Nitrite concentration that completely or almost inhibited AOA ammonia oxidization. The number in parentheses represents the pH of the nitrite concentration inhibition experiment. pH<sup>d</sup>: The range of pH that AOA can grow.

Generally, ammonia tolerance of AOA groups that derived from soils are always higher than the groups from marine and fresh water, and the AOA strains from genus *Ca*. Nitrosocosmicus have the highest currently known tolerance (Table 1). For example, the ammonia tolerance of *Ca*. Nitrosocosmicus franklandus C13 that enriched from arable soil is about 50 times higher than the *Nitrosopumilus maritimus* SCM1 that isolated from tropical marine aquarium, and 2-10 times higher than many soil AOB (Lehtovirta-Morley *et al*., 2016). Because the ammonia tolerant AOA were always undistinguished in many previous studies, archaeal ammonia oxidation and N_2_O production might be underestimated in agricultural soils. Obtaining much more AOA strains from environments and figuring out their biochemical, physiological and ecological features are benefit to uncover the role of AOA playing in global environmental patterns. In our previous work, three AOA strains that affiliating with the genus *Ca*. Nitrosocosmicus were obtained from agricultural soils by using a two-step strategy (Liu *et al*., 2019). In this study, one of these AOA strains that named as *Ca*. Nitrosocosmicus agrestis SS was observed to be able to tolerate 200 mM ammonium (1592.07 μM free ammonia) (Table 1). Based on the physiological and genomic studies, an ammonia tolerant mechanism of the strain *Ca*. Nitrosocosmicus agrestis SS was proposed and verified by using transcriptional gene regulation and metabolites determination, supplying some new features to full the *Nitrosocosmicus* clade or uncover novel roles of AOA in the global environment.

## RESULTS AND DISCUSSION

### Phylogeny and Morphology of “Ca. Nitrosocosmicus agrestis”

Based on the phylogenomic analysis of 43 concatenated universal marker protein sequences, AOA strain SS was found to be closely related to the genus *Ca*. Nitrosocosmicus and form a deep branch in this clade (Fig. 1). The average nucleotide identity (ANI) that calculated between the genomes of strain SS and the other three *Nitrosocosmicus* strains are 72.05%–72.81% (Fig. S1), which are above the proposed genus and below the proposed species boundary thresholds (Varghese *et al*., 2015; Daebeler *et al*., 2018). In accordance with the phylogenomic and genomic ANI analyses, strain SS was assigned to the genus *Ca*. Nitrosocosmicu sand referred to as “*Ca*. Nitrosocosmicus agrestis”.

**Fig. 1.**
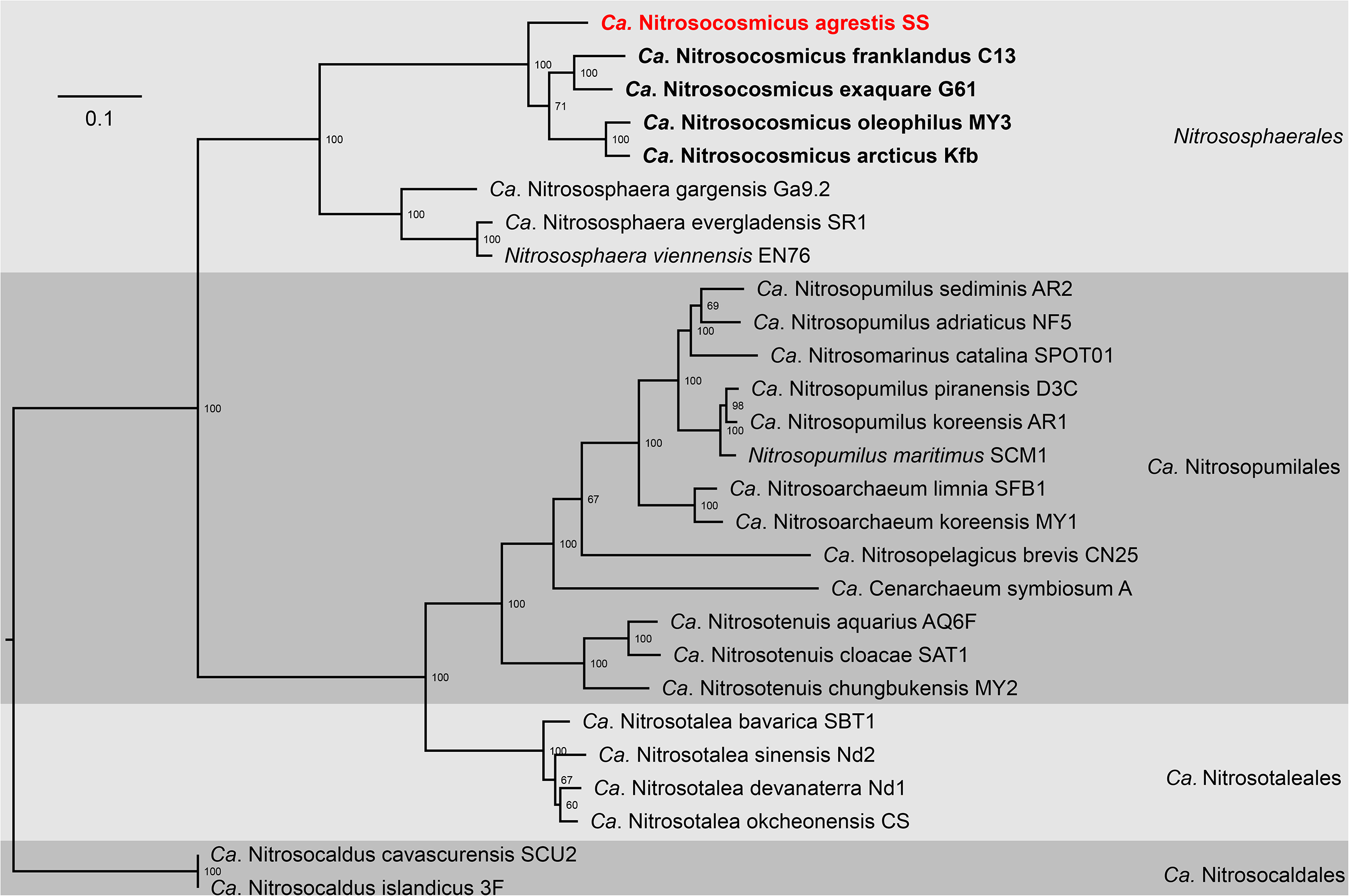
Phylogenomic analysis of “*Candidatus* Nitrosocosmicus agrestis”. The tree was inferred on 43 concatenated universal marker proteins from thaumarchaeota (Table S2) by maximum likelihood with IQ-TREE using a LG+F+R4 model and 1000 replicates.

The TEM and SEM micrographs indicated that these archaeal cells are irregular sphere and 0.8–1.2 μm in diameter (Fig. 2); they often appeared in pairs or aggregation (Fig. 2, Fig. S2). Based on the micrographs, cells of *Ca*. Nitrosocosmicus agrestis were observed to be embedded in a thick extracellular matrix (up to 500 nm), forming an aggregate or biofilm (Fig. 2, Fig. S2). This extracellular matrix could be the putative extracellular polymeric substances (EPS) that secreted during the chemoautotrophic growth of AOA cells. The same feature has also been observed in the cells of *Ca*. Nitrosocosmicus oleophilus MY3 (Jung *et al*., 2016) and *Ca*. Nitrosocosmicus franklandus (Lehtovirta-Morley *et al*., 2016), suggesting the secretion of EPS is probably prevalent in the *Nitrosocosmicus* clade. Even though the flagella that was reported to be essential to the biofilm formation for bacterial cells (Pratt and Kolter, 1998; van Wolferen *et al*., 2018) was not observed on the micrographs (no related gene was identified in the following genomic analysis), the hydrophobic feature of EPS that secreted by cells of *Nitrosocosmicus* clade would facilitate the cells aggregation and adhesion on hydrophobic matrix.

**Fig. 2.**
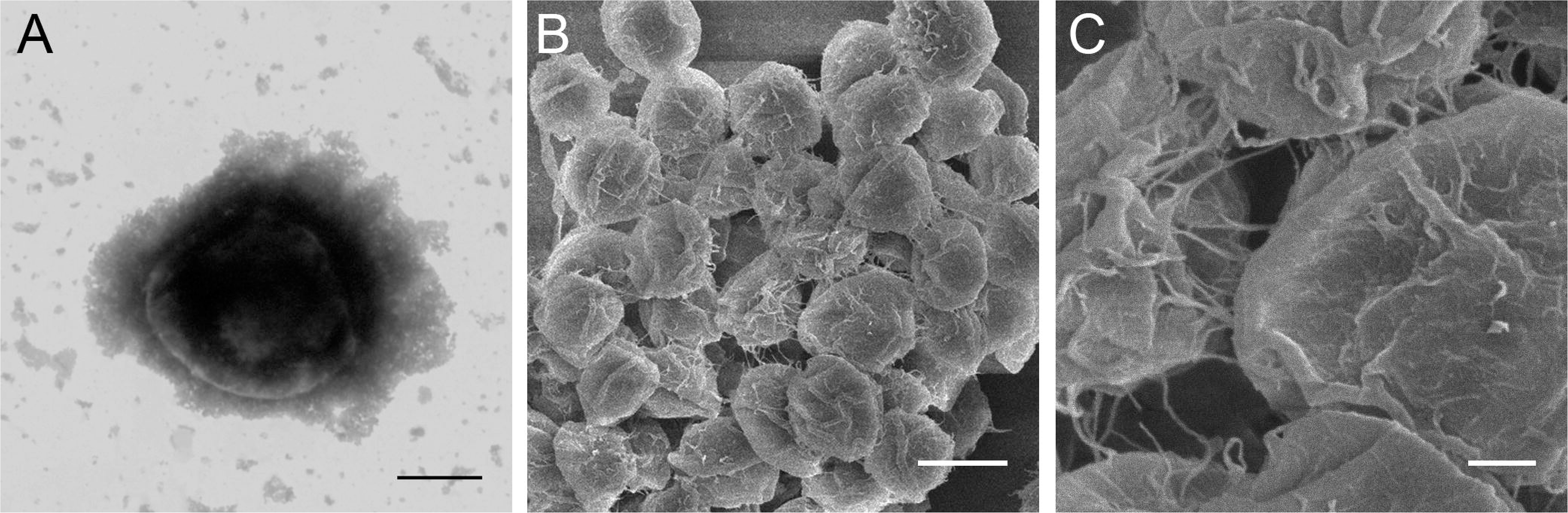
Micrographs of *Ca*. Nitrosocosmicus agrestis cells. (A) TEM observation of *Ca*. Nitrosocosmicus agrestis cell. Scale bar = 500 nm; (B) SEM observation of *Ca*. Nitrosocosmicus agrestis aggregates. Scale bar = 1 μm. (C) SEM observation of *Ca*. Nitrosocosmicus agrestis aggregates. Scale bar = 200 nm.

### Physiology of “Ca. Nitrosocosmicus agrestis”

The ammonia oxidation resulted in the nitrite generation at near-stoichiometric levels (Fig. 3A). A minimum generation time that calculated to be 30.2 h was obtained at 30 °C. This value is shorter than the generation times of strains *Ca*. Nitrosocosmicus exaquare G61 (51.7 h), *Ca*. Nitrosocosmicus franklandus C13 (40 h), and *Ca*. Nitrosocosmicus oleophilus MY3 (77.4 h). After the oxidation of 1 mM substrate (NH_4_^+^), the cell density was determined to be approximately 7.06×10^6^ cells mL ^−1^, which is similar to the levels of *Ca*. Nitrosocosmicus franklandus C13 (7.6×106 cells mL ^−1^) and *Ca*. Nitrosocosmicus oleophilus MY3 (3.2×106 cells mL ^−1^), but lower than the “*Ca*. Nitrosocosmicus exaquare G61” (5.6×107 cells mL ^−1^).

**Fig. 3.**
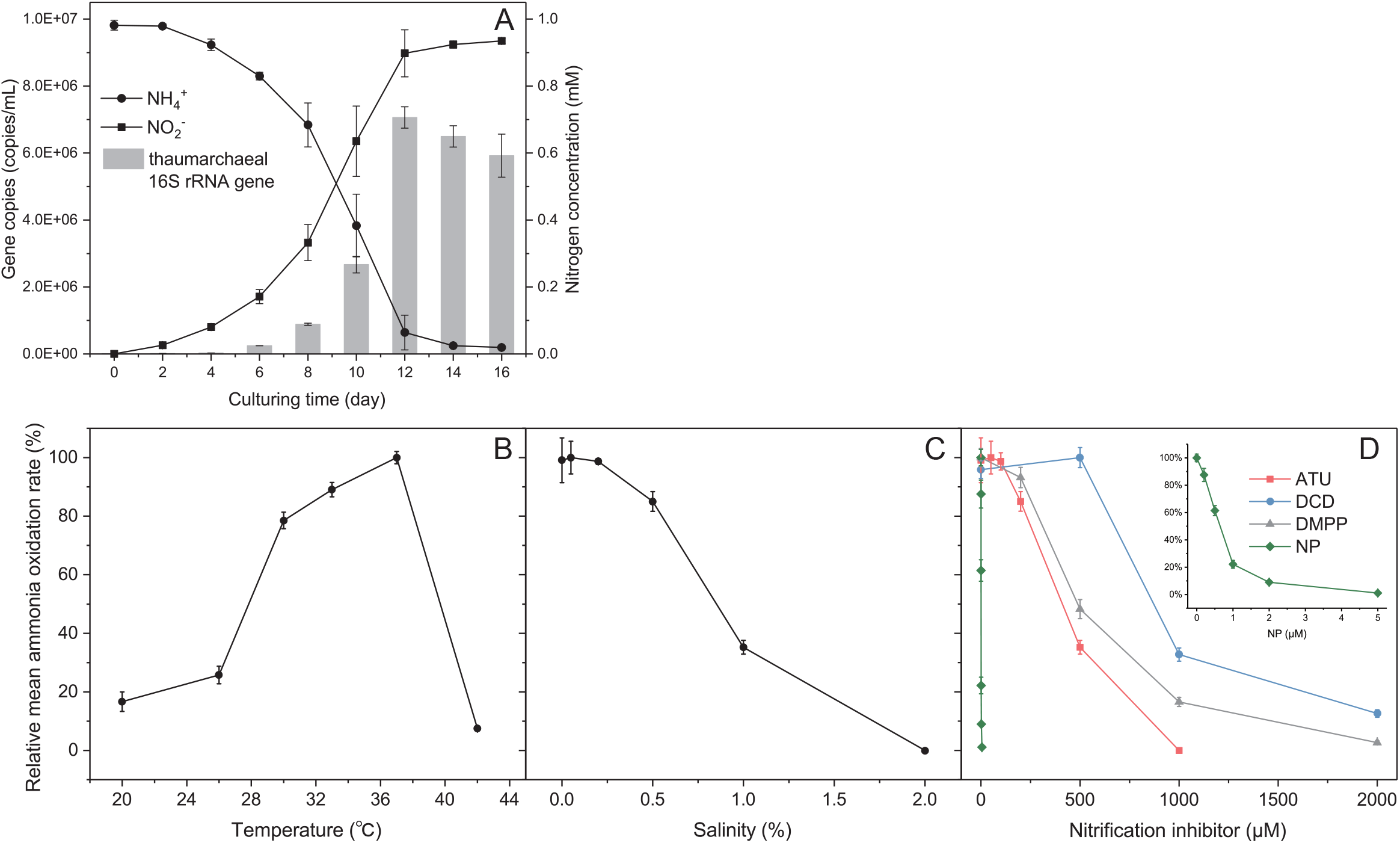
Relationships between ammonia depletion, nitrite production, and archaeal growth in the enrichment culture of strain SS (A), and influences of temperature (B), salinity (C) and nitrification inhibitors (D) on the ammonia oxidation activity of *Ca*. Nitrosocosmicus agrestis. Influence was evaluated by the relative mean ammonia oxidation rates of 8 days cultivation; thaumarchaeotal 16S rRNA genes were measured by qPCR; ATU: allylthiourea; DCD: dicyanodiamide; DMPP: 3,4-dimethylpyrazole phosphate; NP: nitrapyrin; error bars indicate the standard error of the mean for biological triplicates.

*Ca*. Nitrosocosmicus agrestis was able to grow over broad ranges of temperature and salinity. The ammonia-oxidizing activity was observed to be optimal at 37 °C and nearly inhibited at 42 °C (Fig. 3B, Fig. S3); the activity was optimal under the concentration of 0 - 0.2% NaCl and completely suppressed by 2% NaCl (Fig. 3C, Fig. S3). *Ca*. Nitrosocosmicus agrestis was able to tolerate 1% NaCl (kept ~35% of its initial activity), which is much higher than the NaCl tolerance of other terrestrial AOA strains, such as *Ca*. Nitrosotenuis aquarius AQ6f (0.1%) (Sauder *et al*., 2018), *Ca*. Nitrosotenuis cloacae SAT1 (0.03%) (Li *et al*., 2016), *Ca*. Nitrosotenuis uzonensis N4 (0.1%) (Lebedeva *et al*., 2013) and *Ca*. Nitrosoarchaeum koreensis MY1 (0.4%) (Jung *et al*., 2011).

Ammonia oxidation is usually inhibited by the commercial nitrification inhibitors, such as dicyanodiamide (DCD), allylthiourea (ATU), 3,4-dimethylpyrazole phosphate (DMPP) and nitrapyrin (NP); the inhibition varies depending on the type of ammonia oxidizer (Beeckman *et al*., 2018). In the present study, the half-maximal inhibition (IC_50_) of ATU, DCD and DMPP to strain SS were determined to be 445.1, 947.1 and 488.0 μM, respectively (Fig. 3D, Fig. S4, Table 2). These results are similar with the observations in other AOA species (Lehtovirta-Morley *et al*., 2013; Shen *et al*., 2013; Beeckman *et al*., 2018). Differently, the IC_50_ of NP to *Ca*. Nitrosocosmicus agrestis was only 0.599 μM (Table 2), which is far lower than the concentration to *Nitrososphaera viennensis* EN76 (Shen *et al*., 2013), *Ca*. Nitrososphaera sp. JG1 (Kim *et al*., 2012), *Ca*. Nitrosoarchaeum koreensis MY1 (Jung *et al*., 2011) and *Ca*. Nitrosotalea devanaterra (Lehtovirta-Morley *et al*., 2013). NP is a kind of water-insoluble nitrification inhibitor and might have a higher affinity with the hydrophobic surface of *Nitrosocosmicus* (Jung *et al*., 2016) than the other AOA clades. However, *Ca*. Nitrosocosmicus agrestis cannot be extracted by p-xylene like *Ca*. Nitrosocosmicus oleophilus (Jung *et al*., 2016). It may be related to their isolated environment (vegetable soil versus coal tar-contaminated sediment). Therefore, NP has the potential to act as a specific inhibitor of this clade of ammonia oxidizer, but the mechanism is still to be further studied.

**Table 2.** Half-maximal effective concentration [IC_50_ (μM)] of nitrification inhibitors to ammonia oxidation activity of *Ca*. Nitrosocosmicus agrestis

| Nitrification inhibitors | Average IC<sub>50</sub> (μM) | Concentrations tested (μM) |
| --- | --- | --- |
| Allylthiourea (ATU) | 445.1 | 0, 50, 100, 200, 500, 1000 |
| Dicyanodiamide (DCD) | 947.1 | 0, 500, 1000, 2000 |
| 3, 4-Dimethylpyrazole phosphate (DMPP) | 488.0 | 0, 200, 500, 1000, 2000 |
| Nitrapyrin (NP) | 0.599 | 0, 0.2, 0.5, 1, 2, 5 |

The IC<sub>50</sub> were calculated by GraphPad Prism7 refer to shen et al. (Shen *et al.*, 2013).

Although the mechanisms for ammonia tolerance are largely unknown, AOA strains from the genus *Ca*. Nitrosocosmicus were usually reported having higher ammonia tolerance than other AOA (Table 1). In the present study, the ammonia oxidation activity of *Ca*. Nitrosocosmicus agrestis was greatest at 2 mM NH_4_^+^ and decreased with increasing initial ammonium concentration; more than 70% of the initial activity was maintained when the initial ammonium was less than 20 mM (Fig. 4A, B). *Ca*. Nitrosocosmicus agrestis was able to tolerate high concentration of ammonium; at 100 and 200 mM NH_4_^+^, the activities were reduced to approximately 26% and 14% of that at 2 mM NH_4_^+^ (Fig. 4B).

**Fig. 4.**
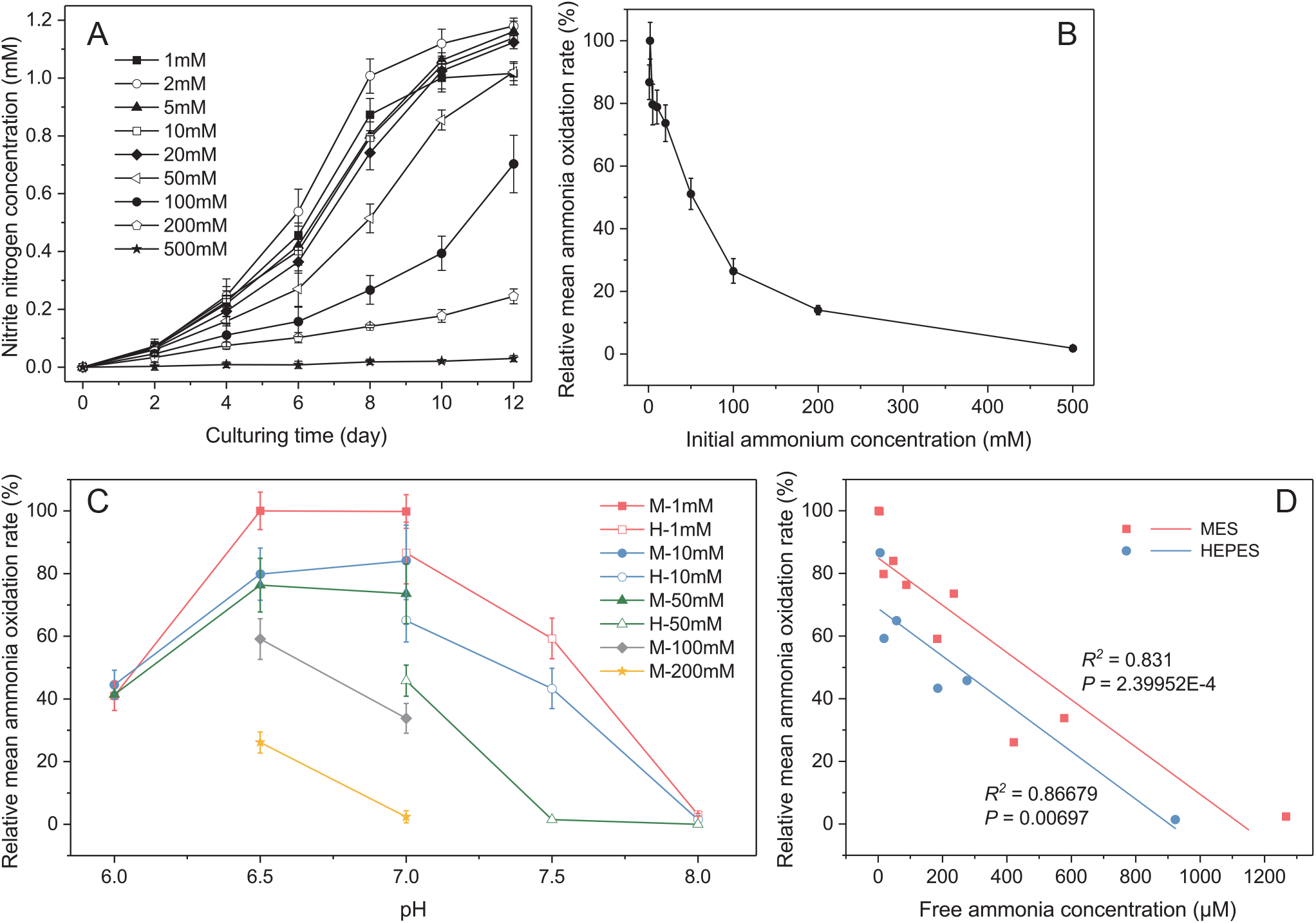
Influences of the initial ammonium (A), nitrite (B), pH (C) and ammonia (D) on the ammonia oxidation activity of “*Ca*. Nitrosocosmicus agrestis”. Influence was evaluated by the relative mean ammonia oxidation rates of 8 days cultivation; The profile of nitrite accumulation with time of ammonia and pH are placed in Fig. S7. (M: MES; H: HEPES; error bars indicate the standard error of the mean for biological triplicates. Free ammonia concentration, calculated according to total ammonia, pH and temperature, estimating by Emerson et al.’s (Emerson *et al*., 1975) formulas.

*Ca*. Nitrosocosmicus agrestis is a neutrophilic microbial strain. Under the low-ammonium concentration (1 and 10 mM), the ammonia-oxidizing activity was detectable during pH range from 6.0 to 7.5, with optimal values being observed in pH 6.5-7.0; the optimal activity was also observed in the range of pH 6.5-7.0 when using 50 mM ammonium as substrate, only the activity was undetectable in pH 7.5 (Fig.4 C, Fig. S6). However, the activities were only detectable in the range of pH 6.5-7.0 when ammonium concentrations were used more than 100 mM. Under these concentrations, the optimal activities were determined in pH 6.5 (Fig.4 C, Fig. S6). Because the higher the pH reaches, the more the free ammonia generates through a pH-dependent equilibrium NH_3_+H^+^ ⇌ NH_4_^+^, which could freely penetrate cell membrane and limit the ammonia oxidation. Then, the more the free ammonia was presented, the lower the ammonia-oxidizing activity was observed; when the ammonia concentration reached 1592.0 μM, ammonia-oxidizing activity was hardly detected (Fig. 4D).

Same as the tolerance to high ammonium, the AOA strains from the genus *Ca*. Nitrosocosmicus were found to have higher nitrite tolerance than the AOA from other genera (Table 1). *Ca*. Nitrosocosmicus agrestis was able to tolerate the nitrite with initial NO_2_^-^ concentration as high as 50 mM; under this concentration, the ammonia-oxidizing activity was reduced to approximately 18% of that at optimal nitrite concentration (Fig. S5). It is obvious that *Ca*. Nitrosocosmicus agrestis has higher nitrite tolerance than the *Nitrosocosmicus* strains *Ca*. Nitrosocosmicus oleophilus (Jung et al., 2016), *Ca*. Nitrosocosmicus franklandus (Lehtovirta-Morley *et al*., 2016), *Ca*. Nitrosocosmicus exaquare (Sauder et al., 2017), as well as the other AOA strains.

Besides the ammonia, *Ca*. Nitrosocosmicus agrestis was able to use urea as sole energy and N source, only the ammonia oxidation rate was one tenth of their growth using ammonia as energy source (Fig. S7). Urea is an abundant form of nitrogen fertilizer in agricultural soils. Many soil AOA strains affiliating with the genera *Nitrososphaera* and *Nitrosocosmicus* have usually been reported to use urea as an energy source (Tourna *et al*., 2011; Jung *et al*., 2016; Lehtovirta-Morley *et al*., 2016; Sauder *et al*., 2017). *Ca*. Nitrosocosmicus franklandus is an ureolytic strain capable of completely converting urea to ammonia and growing on urea as the sole source of ammonia (Lehtovirta-Morley *et al*., 2016). However, the accumulation of ammonia from urea hydrolysis was not observed in *Ca*. Nitrosocosmicus agrestis (Fig. S7B). The ureolytic activity probably varies depending on the type of AOA strain. Besides being used by the soil AOA, urea is suggested to be the energy source for AOA throughout the ocean (Kitzinger *et al*., 2019).

Exogenous supplement of organics were observed to have no significant effect on the ammonia oxidation of *Ca*. Nitrosocosmicus agrestis (Fig. S8). The effect of organic carbon on AOA growth varies between strains. For example, the ammonia oxidation and growth of both *Nitrosotalea* isolates were inhibited by many TCA cycle intermediates but their growth yield increased in the presence of oxaloacetate (Lehtovirta-Morley *et al*., 2014); the pure culture *Nitrososphaera viennensis* requires pyruvate for growth in the laboratory (Tourna *et al*., 2011); the ammonia oxidation of *Ca*. Nitrosocosmicus exaquare that obtained from a municipal WWTP was greatly stimulated by the addition of organics (Sauder *et al*., 2017); α-keto acids could support the growth of some marine AOA via peroxide detoxification (Kim *et al*., 2016). Due to the physiological characters of *Nitrosocosmicus* are largely unknown, the effects of organics on their growth or ammonia oxidation are still needed further profound studies.

### Genome of “Ca. Nitrosocosmicus agrestis”

The genome of *Ca*. Nitrosocosmicus agrestis was recovered by binning from metagenomic assembly (Fig. S9), in which contains 43 contigs with a total length of 3224501 bp. The genome has an average G + C content of 33.42%, contains 3513 protein coding sequences (CDS), and encodes 45 tRNA genes, one 5S rRNA gene, two 16S/23S rRNA operons (Fig. S10). The proportion of coding region of this strain is 70.74%, which is probably the lowest over all AOA strains at present (Table 3).

**Table 3.** Genome features of Ca. Nitrosocosmicus agrestis and other AOA

| AOA strain | Genome size (Mb) | GC (%) | Protein coding genes | coding region (%) | rRNA operons | tRNA genes | Motility | Urease | β-CA | γ-CA-Cam | γ-CA-CamH | Amt-1 | Amt-2 | DME |
| --- | --- | --- | --- | --- | --- | --- | --- | --- | --- | --- | --- | --- | --- | --- |
| *Ca. N. agrestis* SS | 3.22 | 33.41 | 3524 | 70.74 | 2<sup>a</sup> | 45 | - | + | 1 | 1 | 1 | - | 1 | 4 (3)<sup>b</sup> |
| *Ca. N. exaquare* G61 | 2.99 | 33.94 | 3196 | 77.32 | 2 | 36 | - | + | - | 2 | 1 | - | 1 | 4 (3) |
| *Ca. N. oleophilus* MY3 | 3.43 | 34.14 | 3722 | 73.96 | 3 | 38 | - | + | 1 | 3 | 1 | - | 1 | 2 (2) |
| *Ca. N. franklandus* C13 | 2.84 | 34.07 | 3025 | 72.94 | 2 | 39 | - | + | - | 1 | 1 | - | 1 | 3 (3) |
| *Ca. N. arcticus* Kfb | 2.64 | 34.00 | 2970 | 75.83 | 3 | 33 | - | + | 1 | 1<sup>c</sup> | 1 | - | 1 | 3 (1) |
| *N. viennensis* EN76 | 2.53 | 52.72 | 3123 | 87.32 | 1 | 39 | + | + | - | 1 | - | 2 | 1 | 1 |
| *Ca. N. gargensis* Ga9.2 | 2.83 | 48.35 | 3562 | 81.53 | 1 | 37 | + | + | - | - | - | 2 | 1 | 1 |
| *Ca. N. evergladensis* SR1 | 2.95 | 50.14 | 3499 | 83.60 | 1 | 39 | + | + | - | 1 | - | 2 | 1 | 1 |
| *Ca. N. devanaterre* Nd1 | 1.81 | 37.07 | 2067 | 90.99 | 1 | 41 | + | - | - | 1 | - | 1 | 2 | 1 |
| *Ca. N. aquarius* AQ6f | 1.70 | 42.18 | 2008 | 92.90 | 1 | 42 | + | + | - | - | - | 1 | 1 | 1 |
| *Ca. C. symbiosum* A | 2.05 | 57.37 | 2017 | 91.67 | 1 | 45 | - | + | - | - | - | 2 | - | - |
| *Ca. N. brevis* CN25 | 1.23 | 33.16 | 1469 | 94.48 | 1 | 42 | - | - | - | - | - | 1 | 1 | - |
| *Ca. N. limnia* SFB1 | 1.74 | 32.46 | 2038 | 84.83 | 1 | 45 | + | - | - | - | - | 1 | 1 | 2 |
| *Ca. N. catalina* SPOT01 | 1.36 | 31.44 | 1677 | 92.48 | 1 | 43 | - | + | - | - | - | 1 | 1 | - |
| *N. maritimus* SCM1 | 1.65 | 34.17 | 1797 | 90.82 | 1 | 40 | - | - | - | - | - | 1 | 1 | 2 |
| *Ca. N. adriaticus* NF5 | 1.80 | 33.41 | 2206 | 90.61 | 1 | 40 | + | - | - | - | - | 1 | 1 | 1 |
| *Ca. N. piranensis* D3C | 1.71 | 33.82 | 2140 | 90.37 | 1 | 41 | - | + | - | - | - | 1 | 1 | 3 |
| *Ca. N. islandicus* 3F | 1.62 | 41.54 | 1746 | 87.67 | 1 | 39 | + | + | - | - | - | 2 | 1 | - |

b: The number in parentheses indicates the number of copies of the putative polyamine exporter gene.
c: The complete gene sequence cannot be obtained by gene prediction.

*Ca*. Nitrosocosmicus agrestis encodes several genes related to ammonia oxidation, including single copy of the *amoA, amoB* and *amoX*, and three copies of *amoC*. (Table S3). Multiple copies of *amoC* have been reported in the genomes of group I.1b AOA (Tourna *et al*., 2011; Jung *et al*., 2016; Sauder *et al*., 2017); they probably involve in the regulation of stress response to complicated soil environment, as same as they performances in the soil AOB (Berube *et al*., 2007; Berube and Stahl, 2012). A single copy of the gene encoding ammonia transporters (Amt) was also identified in the genome. In general, two types of Amt referring to the high-affinity Amt-1 and the low-affinity Amt-2 were usually found to occur simultaneously in AOA genomes (Offre *et al*., 2014). However, *Ca*. Nitrosocosmicus agrestis and other strains from genus *Ca*. Nitrosocosmicus were found only in the presence of Amt-2 (Table S6, Fig. S11). This is probably closely related to the feature of agricultural soils that the presence of high ammonium; the soil AOA strains, such as the *Nitrosocosmicus-like* are able to rely entirely on Amt-2 to meet the ammonia demand at high level of ammonium.

In addition, the genome contains a cluster of genes for the urea degradation, including the urease subunits, urea accessory proteins and urea transporter (Table S3, S6). These genes were highly expressed under the stimulation of urea (Fig. S7C). The urea degradation supplies ammonia to the ammonia oxidation and could be used as an alternative pathway to produce energy. However, the ammonia oxidation rate of *Ca*. Nitrosocosmicus agrestis on urea was much slower than on ammonia (Fig. S7A). Owing to urease is located in the cytoplasm (predicted by SignalP (Almagro Armenteros *et al*., 2019) and Pred-Signal (Bagos *et al*., 2009)), the ammonia that generated from intracellular urea hydrolysis could be used as N source for the growth but not enough for the energy production via ammonia oxidation. Because of the ammonia in the form of NH_4_^+^ is unable to transport from cytoplasm into periplasm, only through the diffusion in the form of NH_3_, which results in the low substrate concentration for ammonia oxidation that taking place in the periplasmic location (Walker *et al*., 2010; Kozlowski *et al*., 2016). The ureolytic strain *Ca*. Nitrosocosmicus franklandus is able to completely hydrolyze urea to ammonia and grow on urea as the sole source of ammonia (Lehtovirta-Morley *et al*., 2016). However, the mechanism of how urea hydrolysis or ammonia transport in this strain has not yet been given. In addition, the urease gene-containing AOA were observed to dominate the autotrophic ammonia oxidation in acid soils (Lu and Jia, 2013). Expectedly, the mechanism of how urea activates AOA cells remains undefined. It is thus clear that the pathway of urea metabolism fueling ammonia oxidation of AOA would vary between strains and is still far from understood.

Pathways of the central carbon metabolism including CO_2_ fixation, tricarboxylic acid cycle, gluconeogenesis and nonoxidative pentose phosphate pathways were found in the genome of *Ca*. Nitrosocosmicus agrestis and generally similar to those of other AOA (Table S4, Fig. 5). Intriguingly, the genes encoding *β* (D-clades) and γ-classes (Cam and CamH) carbonic anhydrase (CA) were identified in the genome (Fig. S12, Table 3, Table S4,). CA is responsible for the reversible hydration of CO_2_ to HCO_3_^-^ and plays an important role in facilitating carbon transfer into the cell by extracellular converting HCO_3_^-^ to CO_2_, which is subsequently diffuse through the cell membrane (Kerou, Offre, *et al*., 2016). The genes encoding *γ*-class Cam have also been identified in the genomes of *Nitrososphaera viennensis, Ca*. Nitrososphaera evergladensis and *Ca. Nitrosotalea* devanaterra, and genes encoding *α* and *β*-classes CA were usually found in AOB species (Chain *et al*., 2003). In contrast to the location of Cam in periplasmic, the CamH and β-CA of *Ca*. Nitrosocosmicus with no signal peptide are located in cytosolic. Periplasmic CA might constitute an adaption to the discontinuous soils with low CO_2_ availability (Kerou, Offre, *et al*., 2016). The presence of cytosolic CA in the genome suggested that *Ca*. Nitrosocosmicus agrestis has an advantage in advantage in maintaining substrate concentration as well as stabilizing pH in the cytoplasm.

**Fig. 5.**
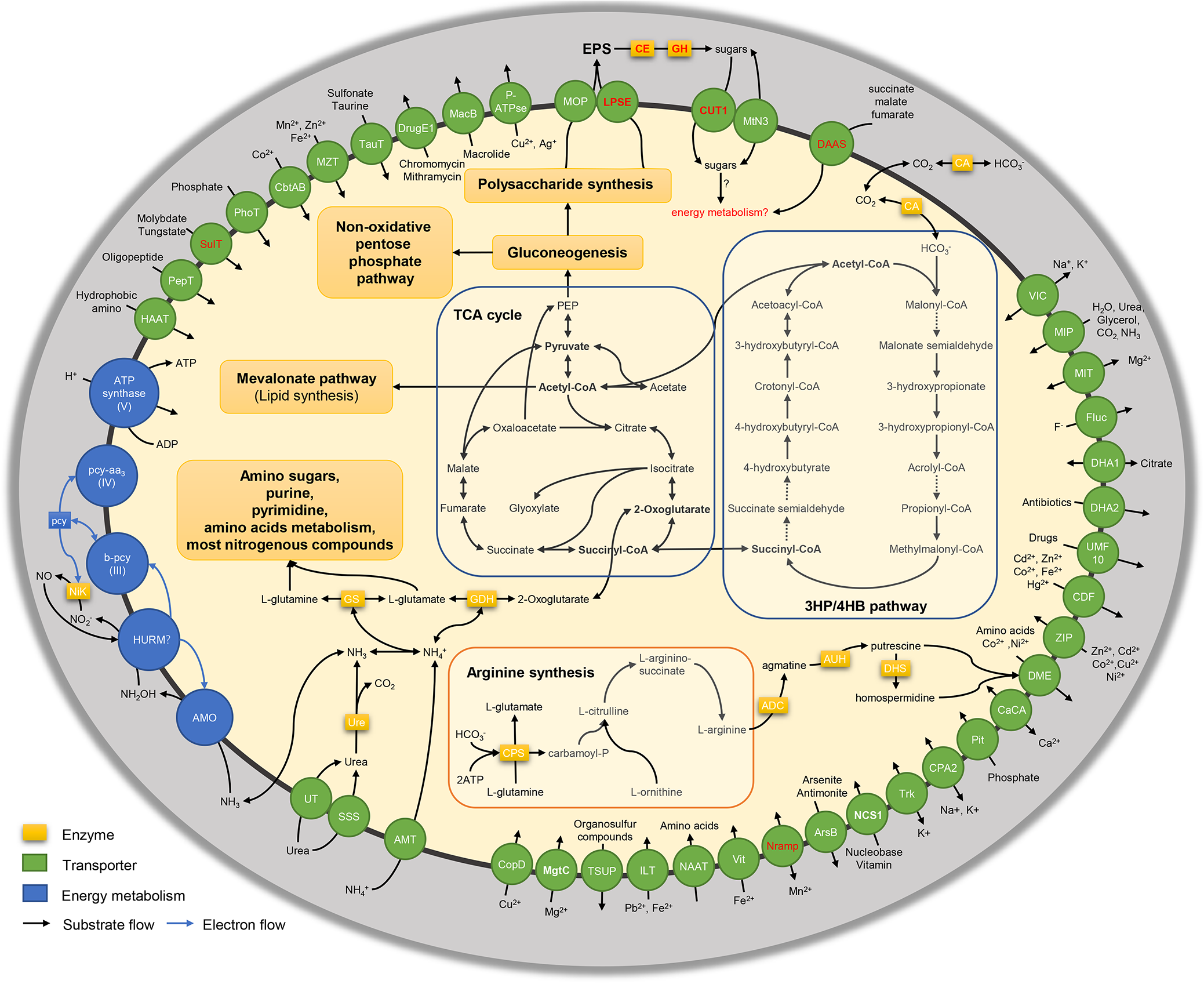
Schematic reconstruction of the predicted metabolic modules and other genome features of *Ca*. Nitrosocosmicus agrestis. Dashed lines indicate reactions for which the enzymes have not been identified. CE: carbohydrate esterase; GH: glycoside hydrolase; CA: carbonic anhydrase; NirK: nitrite reductase; Ure: urease; GDH: glutamate dehydrogenase; GS: glutamine synthetase; ADC: arginine decarboxylase; AUH: agmatinase; DHS: deoxyhypusine synthase. The unique enzymes or transporters of *Nitrosocosmicus* clade are marked in red color. Candidate enzymes, gene accession numbers and transporter classes are list in the supporting information.

A full set of genes encoding for key enzymes involving in the degradation and transport of polysaccharide were found in the genome, including Glycosyl transferases (GTs) family, glycoside hydrolases (GHs) family, carbohydrate esterases (CEs) family, multidrug/oligosaccharidyl-lipid/polysaccharide (MOP) flippase, lipopolysaccharide exporter (LPSE) family, etc (Table S6, S7). These protein families are important to the cell surface modification and EPS production of biofilm-forming bacteria and archaea; an extensive gene encoding these proteins was identified among the AOA groups (such as *N. viennensis, N. gargensis* and *Ca*. Nitrososphaera evergladensis) from Nitrososphaerales (Kerou, Offre, *et al*., 2016). Interestingly, the genes encoding LPSE are found to be unique in the genomes of *Nitrosocosmicus*, indicating the ability to secrete polysaccharide potentially for this AOA clade (Fig. 5). In addition, genes encoding divalent anion/Na^+^ symporter (DASS) and carbohydrate uptake transporter-1 (CUT1) were also found in the genome (Table S6). They catalyze the uptakes of C4-dicarboxylate and complex sugars in bacterial cells (Hugouvieux-Cotte-Pattat *et al*., 2001; Strickler *et al*., 2009). In *Ca*. Nitrosocosmicus agrestis, the sugars derived from the EPS degradation by CEs and GHs are potentially transported into the cells (Fig. 5). However, addition of organic carbons to the cultivation were proved have no effect on the growth or ammonia oxidation of *Ca*. Nitrosocosmicus agrestis (Fig. S8). The uptake of organic carbons (such as the oligosaccharides) by CUT1 is probably not used as part of the energy metabolism but recycled for the EPS production.

### Potential mechanism of ammonia tolerance in “Ca. Nitrosocosmicus agrestis”

All genes encoding the pathway of arginine synthesis were identified in the genome of *Ca*. Nitrosocosmicus agrestis, including carbamoyl phosphate synthetase (CPS), ornithine transcarbamylase (OTC), argininosuccinate synthase (ASS) and argininosuccinate lyase (ASL) (Fig. 5, Table S5). Some genes encoding arginine decarboxylase (ADC), agmatinase (AUH) and deoxyhypusine synthase (DHS) were also found in the genome, which catalyze the conversion of arginine to polyamines, such as putrescine and spermidine (Fig. 5). Meanwhile, four copies of the gene encoding drug/metabolite exporter (DME) were identified in the genome of *Ca*. Nitrosocosmicus agrestis (Fig. 6A). In fact, the DME encoding genes were found in the genomes of almost all of the isolated AOA (Table 3); the genes from genera *Ca*. Nitrosopumilus, *Ca*. Nitrosotalea, *Nitrososphaera, Ca*. Nitrosotenuis, *Ca*. Nitrosoarchaeum and *Ca*. Nitrosocosmicus were clustered into one branch, which is widely distinct from the other branch consisting of *Nitrosocosmicus* and some bacteria and yeast (Fig. 6A). The DME family that existed in these bacteria or yeast were reported be able to mediate the excretion of amino acids and polyamines (Igarashi and Kashiwagi, 2010; Nguyen *et al*., 2015; Lubitz *et al*., 2016), suggesting the active function of DME in AOA strains from genus *Ca*. Nitrosocosmicus. Based on these enzymes and transporters, the NH_4_^+^ in the cytoplasm that derived either from urea hydrolysis or transmembrane diffused NH_3_ could potentially be converted into polyamines and then excreted out by specific transporters, contributing to the high ammonia tolerance of *Ca*. Nitrosocosmicus agrestis.

**Fig. 6.**
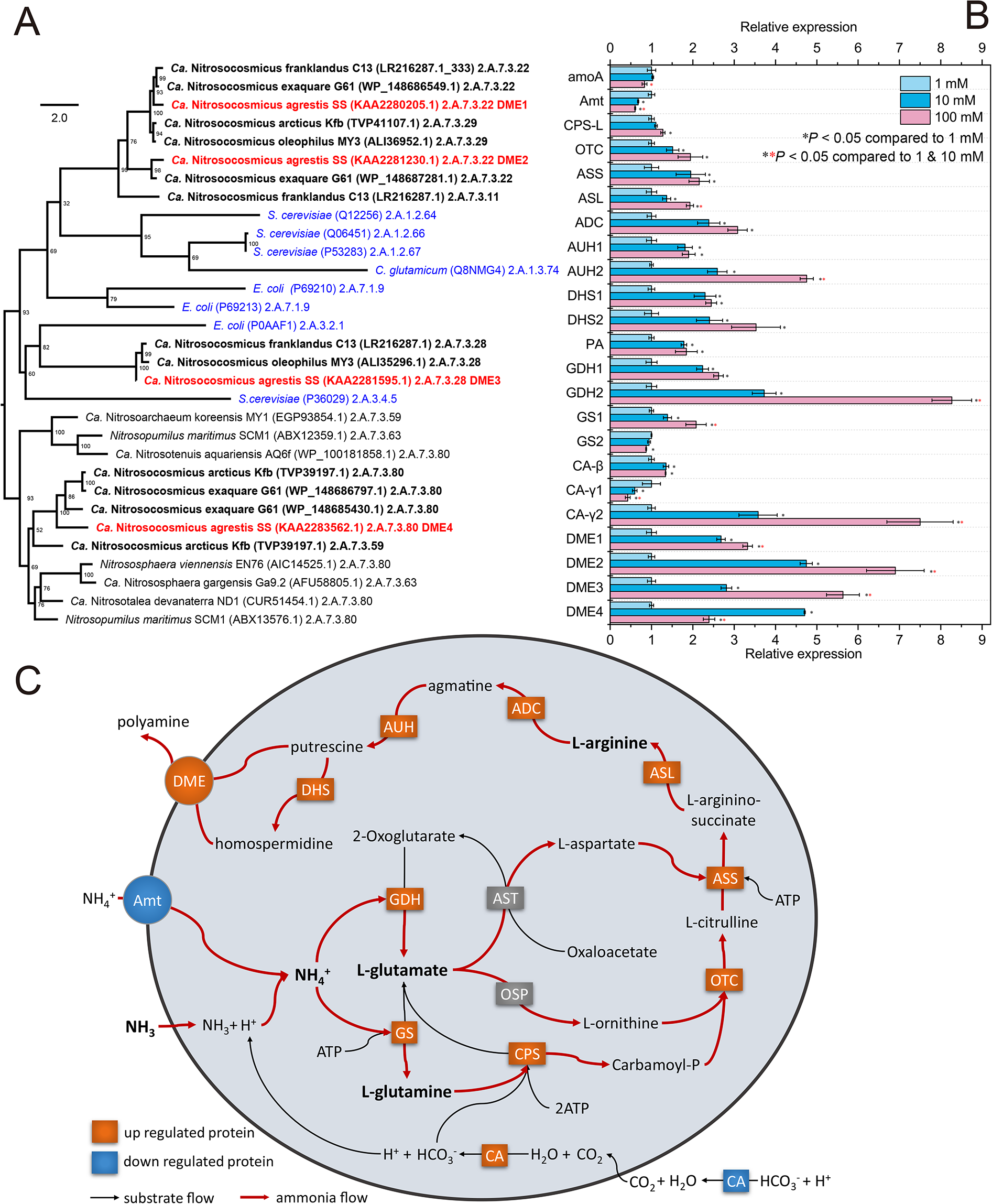
Phylogeny of DME gene (A), relative gene expressions in response to high ammonia (B), and schematic mechanism of ammonia tolerance of Ca. Nitrosocosmicus agrestis (C). The functionally identified DME in bacteria and yeast are marked in blue color; the codes in parentheses are accession number of UniProt or NCBI. *amoA:* ammonia monooxygenase subunit A; Amt: ammonium transporter; CPS: carbamoylphosphate synthase large subunit; OTC: ornithine carbamoyltransferase; ASS: argininosuccinate synthase; ASL: argininosuccinate lyase; ADC: arginine decarboxylase; AUH: agmatinase; DHS: deoxyhypusine synthase; PA: putrescine aminotransferase; GDH: glutamate dehydrogenase; GS: glutamine synthetase; DME: drug/metabolite exporter (DME) family; CA: carbonic anhydrase; error bars indicate the standard error of the mean for technical triplicates; the red arrows represent the flow of ammonia nitrogen.

The relative expressions of genes involved in ammonia oxidation, arginine synthesis, polyamine synthesis, glutamine/glutamate syntheses, carbonic anhydrase, and polyamines transport when responded to high concentration of ammonia were analyzed (Fig. 6B). In comparison with the gene expressions under low ammonia (1 mM NH_4_^+^), the *amoA* had no significant up-regulation and the AMT gene was observed to be down-regulated; however, the arginine synthesis, polyamine synthesis and putative DME polyamine exporter related genes were observed to have up-regulation with the increase of ammonia concentration. In addition, the gene encoding putrescine aminotransferase (PA) that catalyzes the first step of putrescine degradation was also up-regulated in response to high ammonia, suggesting the more production of putrescine under high ammonia. The glutamate dehydrogenase (GDH) and glutamine synthetase (GS) genes were also found to be significantly up-regulated, which depleted cytoplasmic ammonium and resulted in the production of glutamine, providing more precursors for ornithine synthesis. Up-regulation of carbonic anhydrase (CA) genes that locating in the intracellular (β-CA and CamH of γ-CA) can provide H^+^ to avoid cytoplasmic pH rise caused by ammonia uptake and provide more bicarbonate for carbamoylphosphate synthesis. In summary, the up-regulations of genes involving in polymaines synthesis and secretion when in response to high ammonia were probably responsible for the high ammonia tolerance of *Ca*. Nitrosocosmicus agrestis. (Fig. 6C).

To prove the above proposed mechanism, the energy utilization and the polymaines production were studied. In contrast to the ammonia oxidation in low substrate (1 mM), the average ammonia oxidation rate of *Ca*. Nitrosocosmicus agrestis exposing to high substrate (100 mM) was decreased to 83% (Fig. 7A); if these energy that derived from ammonia oxidation were completely converted into biomass (determined by protein concentration), the energy utilization efficiency was only 10% (Fig. 7B). It is suggested that a large amount of energy was used for some other syntheses in order to tolerant high ammonia, such as the polyamines synthesis. Based on the detection by HPLC-MS/MS, the polyamines putrescine, cadaverine and spermidine were identified from the supernatant, with concentrations ranging from 1.37 to 10.25 μg L^−1^ (Fig. 7C). In contrast to the productions under low ammonium, the cadaverine concentration had no difference but the spermidine concentration was about three times higher and the putrescine (10.25 μg L^−1^) was only detected in high amm onium. In the actual situation, the polyamines would have higher concentrations due to (i) the presence of other type of polyamines, (ii) the absorption by EPS for they have positive charge, and (iii) the uptake by bacterial partners. The polyamines are kinds of dissolved organic carbon (DOC), which could be served as carbon source for the growth of heterotrophic bacteria when being released into ambient environment. Recently, some marine AOA *Nitrosopumilus* spp. were reported to be able to release hydrophobic amino acid through passive diffusion and then fuel the prokaryotic heterotrophy in the ocean (Bayer *et al*., 2019). Though very few amino acid was detected in the supernatants neither from low ammonium culture nor from high ammonium culture (Fig. S13), the released polyamines were still able to fuel the heterotrophic partners. In fact, during the cultivation of *Ca*. Nitrosocosmicus agrestis, the utilization of high ammonium as substrate has undoubtedly resulted in lower abundance of AOA in comparison with the low ammonium culture (data not shown), suggesting the promotion of bacterial growth by high ammonium stress. Further studies are needed to figure out if the polyamines that secreted under high ammonium are responsible for the interaction between AOA and heterotrophy, which subsequently hinder the AOA enrichment and even isolation.

**Fig. 7.**
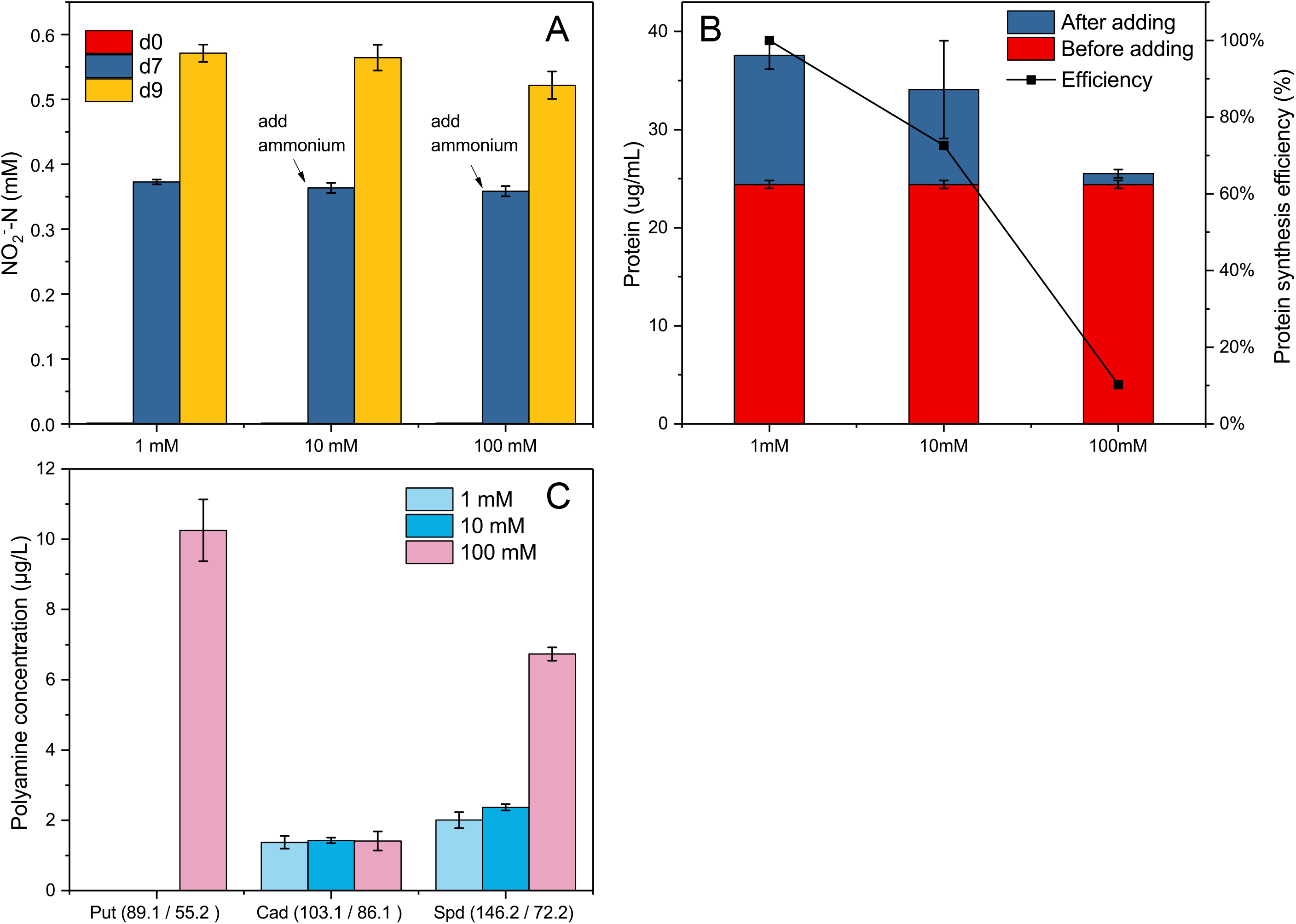
**After adding different concentrations of ammonia nitrogen for two days, the** accumulation of nitrite (A), protein synthesis (B), and accumulation of polyamine (C) by *Ca*. Nitrosocosmicus agrestis. Put: putrescine; Cad: cadaverine; Spd: spermidine; error bars indicate the standard error of the mean for biological triplicates.

## CONCLUSION

Physiological characterization of *Ca*. Nitrosocosmicus agrestis has revealed its extensive environmental adaptability, especial for the high ammonia tolerance. Genomic analysis indicated that *Ca*. Nitrosocosmicus agrestis harbors a variety of carbohydrate-active enzymes and transporters relating to the EPS synthesis, which is potentially responsible for they adaption to the complex environment. Surprisely, *Ca*. Nitrosocosmicus agrestis has a complete pathway for the polyamines synthesis and secretion in genome, suggesting a mechanism to tolerate high ammonia. High ammonium resulted in the higher productions of putrescine and spermidine, proving the polyamine synthesis is the adaption strategy in response to high ammonia.

Strain SS is an ammonia oxidizing archaeon enriched from soil, we propose the following provisional taxonomic assignment:

*Nitrososphaerales* order

*Nitrososphaeraceae* fam. and

*Candidatus* Nitrosocosmicus agrestis sp. nov.

Etymology: nitrosus (Latin masculine adjective), nitrous; cosmicus (Latin masculine adjective): cosmopolitan; agrestis (Latin masculine adjective), belonging to the field.

Locality: vegetable field in Suishi village, Guangzhou, Guangdong, China

Diagnosis: an ammonia oxidizer growing chemolithoautotrophic, appearing as 0.8-1.2 μm irregular cocci covered by putative exopolysaccharides, optimum pH 6.5-7.0, optimum temperature 37 °C, optimum salinity 0-0.2%, utilize urea, ammonia oxidation activity can still be detected at a free ammonia concentration of 1592 μM but almost completely inhibited by 2 μM nitrapyrin.

## MATERIALS AND METHODS

### Culture maintenance

*Ca*. Nitrosocosmicus agrestis was obtained from a vegetable field after the enrichment using a two-step strategy (Liu *et al*., 2019). The AOA abundance in the enrichment culture was ≥ 95% (symbiotic bacteria mainly belonging to *Rhizobiales* (Fig. S9)).

*Ca*. Nitrosocosmicus agrestis was incubated at 30 °C in the dark without shaking, maintaining in a mineral salts medium (MSM) containing (per litre): 1 g NaCl, 0.4 g MgCl_2_·6H_2_O, 0.1 g CaCl_2_·2H_2_O, 0.2 g KH_2_PO_4_, 0.5 g KCl, 1 mL Fe-EDTA solution (49.14 g L^−1^ FeSO_4_·7H_2_O and 196.896 g L^−1^ EDTA disodium salt dihydrate), 2.5 mL NaHCO_3_ solution (1 M) and 1 mL NH_4_Cl solution (1M), 1mL trace element solution (1.5 g L^−1^ FeCl_2_·4H_2_O, 190 mg L^−1^ CoCl_2_·6H_2_O, 100 mg L^−1^ MnCl_2_·6H_2_O, 70 mg L^−1^ ZnCl_2_, 62 mg L ^−1^ H_3_BO_3_, 36 mg L^−1^ Na_2_MoO_4_·2H_2_O, 24 mg L^−1^ NiCl_2_·6H_2_O, and 17 mg L^−1^ CuCl_2_·2H_2_O). Solutions were filtered using 0.22 μm sterilized filters. To eliminate symbiotic bacteria, an antibiotic mixture containing with ciprofloxacin (50mg/L) and azithromycin (50mg/L) was added. The initial pH value was adjusted to 7.0. Ten percent (w/v) of quartz (1 mm diameter) was supplied as the attachment of archaeal cells during the seed cultivation and transferred into a fresh medium.

### Physiological characterisation

To prepare AOA culture for physiological characterisation, the quartzs in seed culture were collected and the AOA cells attaching on quartz were washed out by using MSM; 10% eluate was inoculated into the fresh MSM without supplying quartz. The effects (tolerances) of ammonium concentration were determined in MSM added with 1-500 mM NH_4_Cl and inhibition by nitrite concentration by supplementation with 0-100 mM NaNO_2_. The effects of temperature, initial pH and salinity on ammonia oxidation were also investigated with values in the range of 20-42 °C, 6.0-8.0 and 0-2.0%, respectively. To avoid the rapid pH decrease due to bicarbonate uptake and ammonia oxidation, the MSM that used for pH characterization was not supplied with NaHCO_3_ (the diffused CO_2_ is sufficient for SS growth) and buffered with 10mM MES (2-(N-morpholino)ethanesulfonic acid), HEPES (4-(2-hydroxyethyl)-1-piperazineethanesulfonic acid), Tris (Tris(hydroxymethyl)aminomethane) between the range of pH 6-7, 7-8, and 8-9, respectively. The effects of organics on ammonia oxidation were determined in MSM supplemented with 50 mg/L each organic carbon.

All physiological characteristics were determined by evaluating the ammonia oxidation activity during batch cultivation. Concentrations of nitrite and ammonium were determined by standard Griess-Ilosvay method and indophenol blue method, respectively (ISO/TS 14256–1:2003, 2003). The mean ammonia oxidation rate was calculated basing on the determination of nitrite concentration.

The density of AOA in the enrichment was analyzed using absolute quantitative PCR based on 16S rRNA gene primers (SS16S-1F / SS16S-1R), following the method described in the previous study (Liu *et al*., 2019).

### Microscopy

To prepare samples for the transmission electron microscopy (TEM), AOA cells were collected by the centrifugation at 8000 *g* for 10 min and fixed with 2.5% glutaraldehyde solution at 4 °C for 12 h. After being washed in PBS buffer for 3 times, the cells were fixed with 1% osmic acid solution for 2 h. After being washed in PBS buffer for 2 times, the samples were dehydrated in a graded series of ethanol solutions (50%, 70%, 90%, 100%) at 4 °C for 10 min and transferred onto a copper grid. Each preparation was observed with H-7650 transmission electron microscope (Hitachi, Tokyo, Japan) at an accelerating voltage of 80 kV.

To prepare samples for scanning electron microscopy (SEM), glass slides (1 cm^2^) were placed in the bottom of flask and used as the attachment of AOA cells during cultivation. After the incubation, the slides were directly used for the fixation with glutaraldehyde, the dehydration by a graded series of 100% absolute ethanol, and the critical point dry using an Autosamdri-815 series A (Tousimis, Baltimore, USA). The slides were mounted on an aluminum stub and sputter-coated with platinum using a sputter coater EMS150T (Quorum, East Sussex, UK), then imaged using Merlin field emission scanning electron Microscope (Carl Zeiss, Jena, Germany).

### Genome sequencing, assembly and annotation

The genomic DNA extraction from 3,000 mL batch cultures was performed according to a previously described protocol (Liu *et al*., 2019). The purified DNA was fragmented to ~400 bp with the aid of an M220 focused-ultrasonicator (Covaris, Woburn, USA) and subsequently used for library preparation using a TruSeq™ DNA Sample Prep Kit (Illumina, San Diego, USA). Metagenomic sequencing was performed on a HiSeq X Ten sequencing system (Illumina, San Diego, USA).

The raw data was length- and quality-filtered using Trimmomatic v0.38 (Bolger *et al*., 2014) and de novo assembled using metaSPAdes v3.13.0 (Nurk *et al*., 2017) with 127-mer. Then, GapCloser module of SOAPdenovo2 v1.12 (Luo *et al*., 2012) was used with default parameters to fill a proportion of gaps of the assembled scaffolds. Genome binning of the metagenomic assemblies was conducted using MaxBin2 v2.26 (Wu *et al*., 2016) with the universal marker gene sets (Wu *et al*., 2013). Scaffolds with length <1000 bp in assembly were removed from the further analysis. The completeness, contamination and strain heterogeneity of genome were evaluated using CheckM v1.013 (Parks *et al*., 2015). Gene prediction and annotation were done using JGI-IMG/MER (Chen *et al*., 2019) and MicroScope platform (Vallenet *et al*., 2019). KEGG pathways were generated using KASS (Moriya *et al*., 2007). Putative transport proteins and carbohydrate-active enzymes were identified using TCDB (Saier *et al*., 2014) and dbCAN2 (Zhang *et al*., 2018), respectively. The average nucleotide identity (ANI) of four *Nitrosocosmicus* genome were calculated using JSpeciesWS (Richter *et al*., 2016). The genome described in this study has been deposited in NCBI under GenBank accession VUYS00000000.

### Phylogenomic analysis

A phylogenomic tree was obtained using the automatically generated alignment of 43 concatenated universal marker protein sequences (Table S2), which were identified by CheckM v1.013 (Parks *et al*., 2015). The best-fit model of evolution was selected with ModelFinder (Kalyaanamoorthy *et al*., 2017) and phylogenomic trees were inferred by Maximum-likelihood with IQ-TREE v1.6.1 (L. T. Nguyen *et al*., 2015).

### Relative expression of genes

To extract total RNA from liquid culture, archaeal cells in the enrichments were collected from biological triplicates by filtering through a 0.22μm cellulose filter; the filter was cut into pieces and placed in a grinding tube containing 0.5 g quartz sand. RNA extraction was performed using the RNAiso Plus (Takara, Dalian, China), and the first strand of cDNA synthesis was performed using the PrimeScript™ RT reagent Kit with gDNA Eraser (Takara, Dalian, China). All quantitative PCR were performed in triplicate on an ABI 7500 Fast real-time PCR system (Applied Biosystems, Carlsbad, USA) using TransStart Tip Green qPCR SuperMix (Transgen, Beijing, China). The reaction condition was as follows: 1 min at 94 °C; 40 cycles of 10 s at 94 °C and 34 s at 60 °C. The 2^−ΔΔCt^ method was used to estimate fold change in gene expression, normalized to the endogenous control 16S rRNA. Primer sets (Table S9) were designed on the function genes from *Ca*. Nitrosocosmicus agrestis genome.

### Determination of protein content

To measure the total protein in AOA cells, about 20 mL of enrichment culture was collected and filtered through a 0.22 μm cellulose filter; the filter retaining with cells was collected and cut into pieces and placed in a 2 mL grinding tube containing 0.5 g quartz sand. 500 μL of lysis buffer (TE buffer containing 1% Triton X100) was added into the tube and mixed for 5 min using the Vortex Adapter (13000-V1-24, QIAGEN, Germany). After the centrifugation at 5,000 g for 5 min, the pellet was collected and used for the protein measurement by BCA Protein Assay Kit (Sangon Biotech, Shanghai, China).

### Quantification of polyamines and amino acids in supernatant

After the centrifugation, 800 μL of the supernatant was collected and filtered through a 0.22 μm cellulose filter. The filtrate was mixed with 200 μL of sulfosalicylic acid (10%) and incubated at 4 °C for 1 hour, following a centrifugation at 14,000 g for 15 min. The supernatant was filtered through a 0.22 μm cellulose filter again and used for the quantitative analysis of amino acids or polyamines.

Using LC-20A HPLC system (shimadzu, Japan) coupled with API 4000 Quantiva triple quadrupole tandem mass spectrometer (SCIEX, USA) to analyze putrescine, cadaverine, spermidine, and spermine in the supernatant (HPLC–MS/MS experimental conditions of each polyamine were listed in Table S10) (Gosetti *et al*., 2013). HPLC separation was performed at 0.30 mL/min and the column compartment to 40 °C. The stationary phase was an Inertsil^®^ ODS-3 column (2.1×100 mm, 3 μm), and the mobile phase was an isocratic mixture (A: B=20:80) of 5 mmol/L CH_3_COONH_4_ in H_2_O (A) and 0.2% HCOOH in H_2_O (v/v) (B). Using amino acid analyser A300-advanced (membraPure, Berlin, Germany) to analyze seventeen dissolved amino acids (Cys, Asp, Met, Thr, Ser, Glu, Pro, Gly, Ala, Val, Ile, Leu, Tyr, Phe, His, Lys, Arg) in the supernatant.

## Compliance with ethical standards

Conflict of interest: The authors declare that they have no conflict of interest.

---

**FIGURE S1** Heat maps showing pairwise ANI values inferred from the available AOA genomes.

**FIGURE S2** Transmission electron micrograph of *Ca*. Nitrosocosmicus agrestis cells in pairs (A) (scale bar = 500 nm) and aggregates (B) (scale bar = 2 μm). (C) Scanning electron micrograph of strain SS cells incubating with 50mg/L ciprofloxacin and azithromycin. Scale bar = 500nm.

**FIGURE S3** Effects of temperature (A) and salinity (B) on the ammonia oxidation activity of *Ca*. Nitrosocosmicus agrestis. Error bars indicate the standard error of the mean for biological triplicates.

**FIGURE S4** Effects of nitrification inhibitors ATU (A), DCD (B), DMPP (C) and NP (D) on the ammonia oxidation activity of *Ca*. Nitrosocosmicus agrestis. Error bars indicate the standard error of the mean for biological triplicates.

**FIGURE S5** (A)Variation of nitrate nitrogen concentration with time at different initial nitrite concentration; (B) the average ammonia oxidation rate of homogenization at different initial nitrite concentration. Error bars indicate the standard error of the mean for biological triplicates.

**FIGURE S6** Effect of 1mM (A), 10mM (B), 50mM(C), 100mM (D) and 200mM (D) concentration of ammonium on ammonia oxidation activity of strain SS at different pH. Error bars indicate the standard error of the mean for biological triplicates.

**FIGURE S7** Effect of urea on nitrite (A) or ammonia (B) concentration in ammonia oxidation process, and relative gene expressions in response to urea (C). 1A, 0.1A, 0A: 1mM, 0.1mM, 0mM NH_4_Cl; 1U: 1mM urea. Amt: Amt ammonium transporter; UreC: urease subunit alpha; UT: urea transporter. Error bars indicate the standard error of the mean for biological triplicates.

**FIGURE S8** Growth of *Ca*. Nitrosocosmicus agrestis amended with organic carbon and incubated with ciprofloxacin-azithromycin (A) or no antibiotics (B). The initial ammonia concentration is 1mM. Error bars indicate the standard error of the mean for biological triplicates.

**FIGURE S9** Classification and coverage of metagenome contigs (A) and composition of bins from the *Ca*. Nitrosocosmicus agrestis cultures (B); bin2 is the genome of *Ca*. Nitrosocosmicus agrestis.

**FIGURE S10** Gene arrangement around 16S/23S operons in the genomes of *Ca*. Nitrosocosmicus. SS: *Ca*. Nitrosocosmicus agrestis SS; G61: *Ca*. Nitrosocosmicus exaquare G61; MY3: *Ca*. Nitrosocosmicus oleophilus MY3.

**FIGURE S11** Evolution Analysis of Amt amino acid sequences in the different AOA genomes. Software: iqtree; Method: maximum likelihood; Model: LG+F+R3; bootstrap value: 1000.

**FIGURE S12** Evolution analysis of β (A) or γ (B)-carbonic anhydrase based on amino acid sequence. Software: iqtree; Method: maximum likelihood; Model: LG+I+G4; bootstrap value: 1000.

**FIGURE S13** Detection of dissolved free amino acid in the supernatant of the enrichments under different concentrations of ammonium. Except for (Cys)2 of 50 nmol / mL, all other standards are 100 nmol / mL.

**TABLE S1** Evaluation of genomes by CheckM software.

**TABLE S2** 43 concatenated universal marker proteins for phylogenomic analysis.

**TABLE S3** Genes of energy metabolisms in *Ca*. N. agrestis genome.

**TABLE S4** Genes of carbon metabolisms in *Ca*. N. agrestis genome.

**TABLE S5** Genes of nitrogen metabolisms in *Ca*. N. agrestis genome.

**TABLE S6** Genes of transporters in *Ca*. N. agrestis genome.

**TABLE S7** Genes of carbohydrate-active enzymes in *Ca*. N. agrestis genome.

**TABLE S8** DME family transporters of AOA and polyamine exporters

**TABLE S9** Primers used in this study

**TABLE S10** HPLC–MS/MS experimental conditions of polyamines

