## Supplementary material for "Physiological and genomic analysis of “*Candidatus* Nitrosocosmicus agrestis”, an ammonia tolerant ammonia-oxidizing archaeon from vegetable soil": Figure S1-S13

**TABLE S1** Evaluation of genomes by CheckM software.

**TABLE S2** 43 concatenated universal marker proteins for phylogenomic analysis.

**TABLE S3** Genes of energy metabolisms in *Ca.* N. agrestis genome.

**TABLE S4** Genes of carbon metabolisms in *Ca.* N. agrestis genome.

**TABLE S5** Genes of nitrogen metabolisms in *Ca.* N. agrestis genome.

**TABLE S6** Genes of transporters in *Ca.* N. agrestis genome.

**TABLE S7** Genes of carbohydrate-active enzymes in *Ca.* N. agrestis genome.

**TABLE S8** DME family transporters of AOA and polyamine exporters

**TABLE S9** Primers used in this study

**TABLE S10** HPLC–MS/MS experimental conditions of polyamines

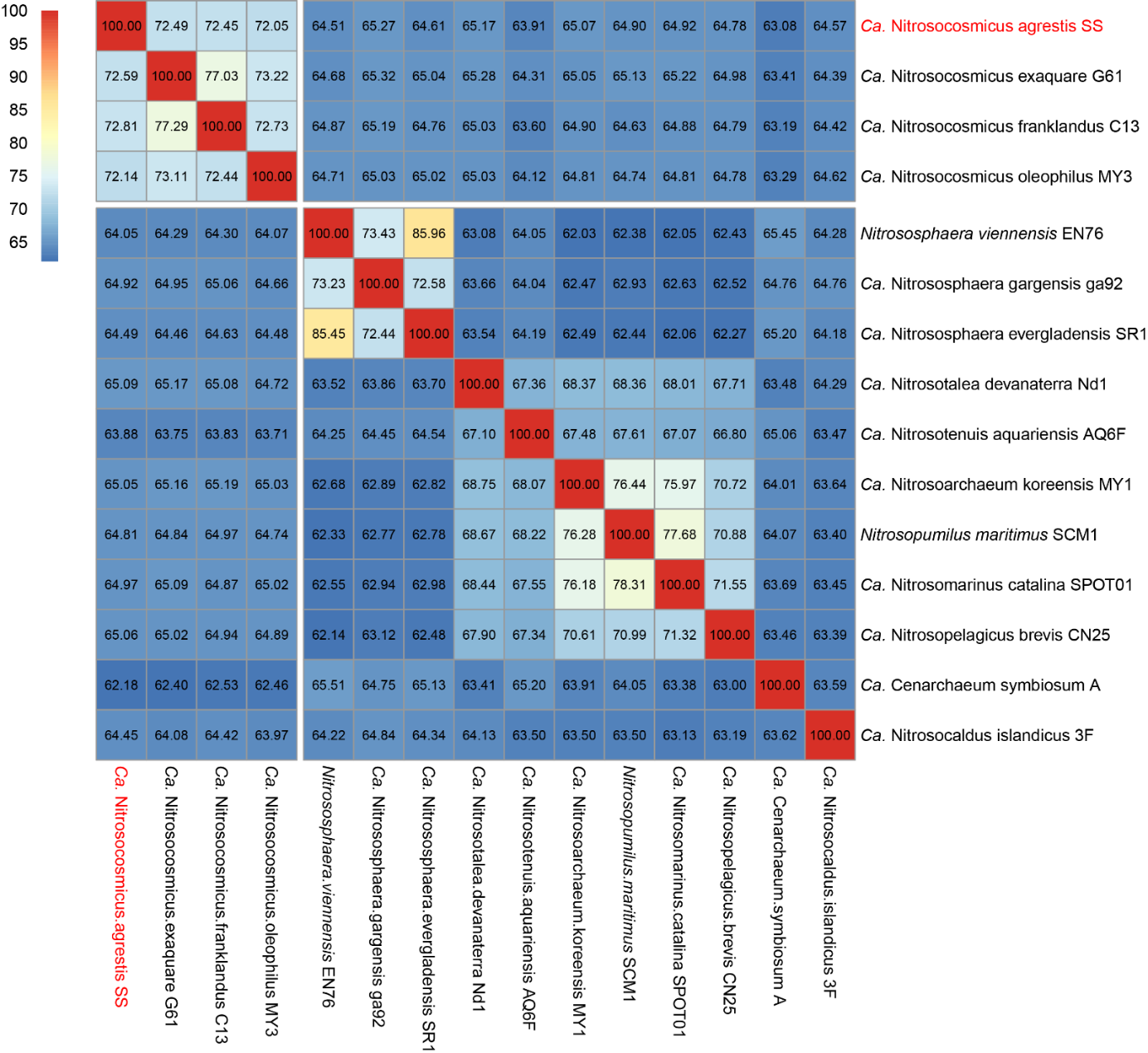

**FIGURE S1** Heat maps showing pairwise ANI values inferred from the available AOA genomes.

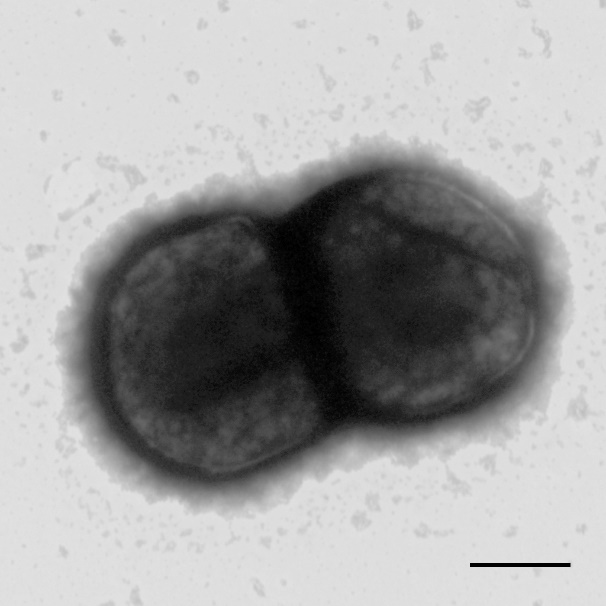

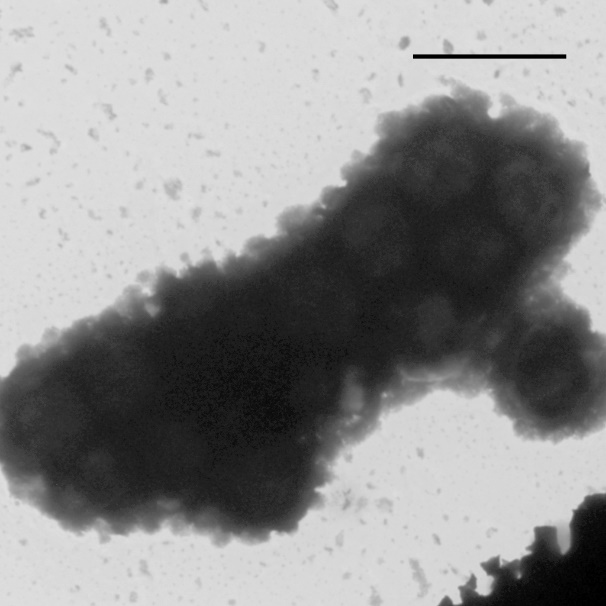

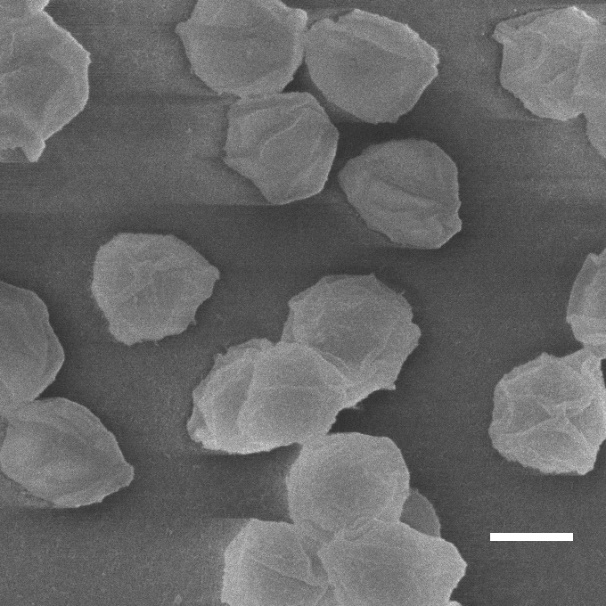

**C**

**B**

**A**

**FIGURE S2** Transmission electron micrograph of *Ca.* Nitrosocosmicus agrestis cells in pairs (A) (scale bar = 500 nm) and aggregates (B) (scale bar = 2 µm). (C) Scanning electron micrograph of strain SS cells incubating with 50mg/L ciprofloxacin and azithromycin. Scale bar = 500nm.

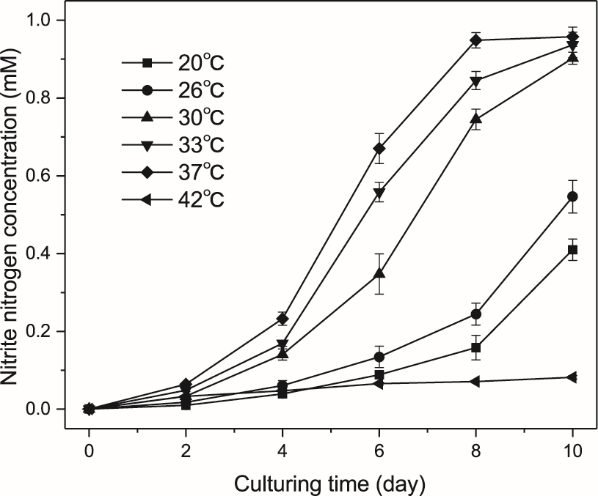

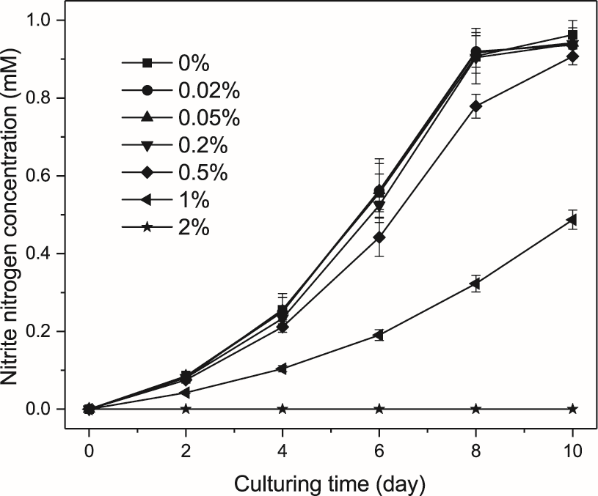

**B**

**A**

**FIGURE S3** Effects of temperature (A) and salinity (B) on the ammonia oxidation activity of *Ca.* Nitrosocosmicus agrestis. Error bars indicate the standard error of the mean for biological triplicates.

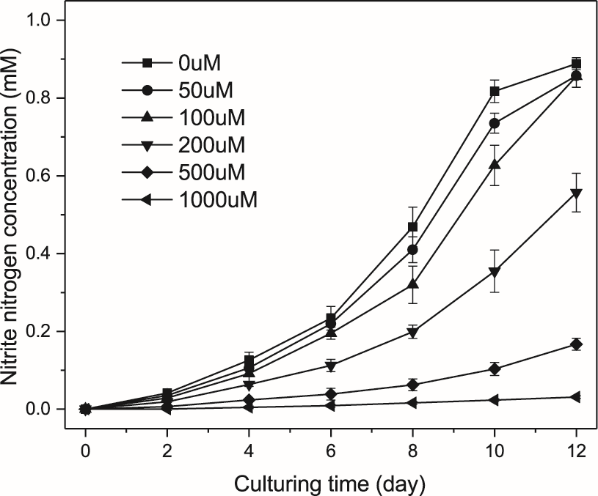

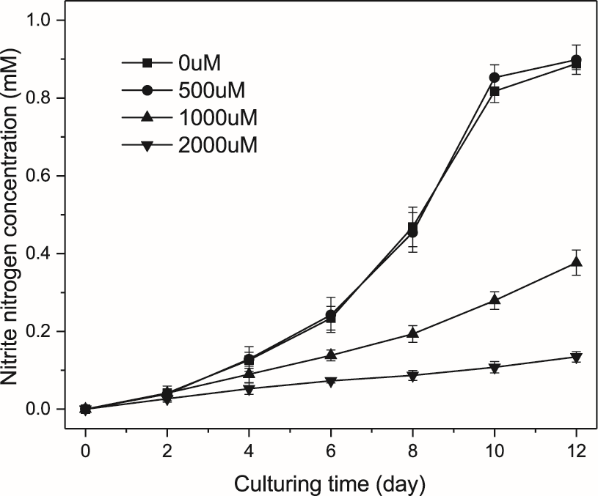

**B**

**A**

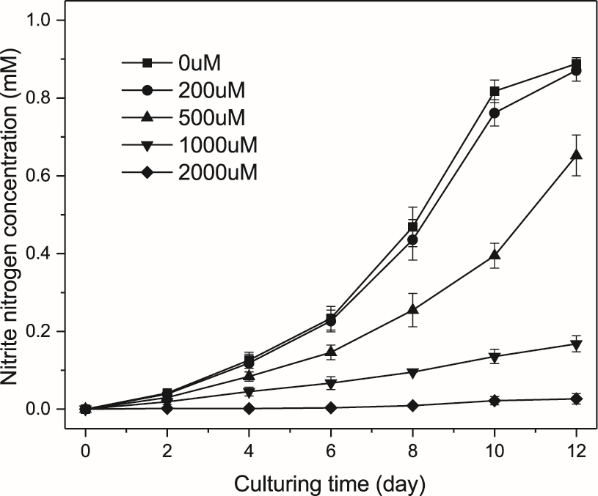

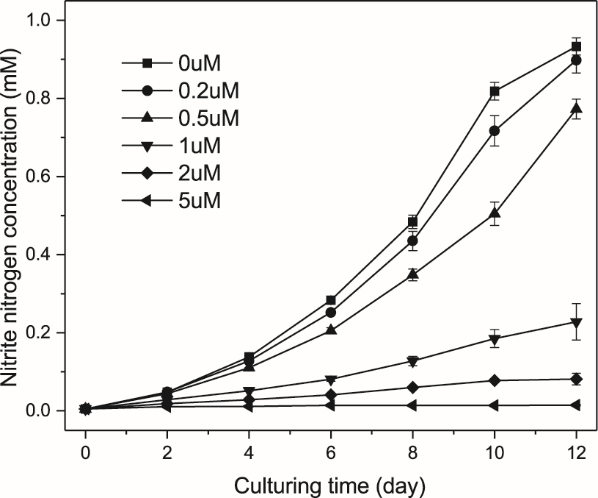

**C**

**D**

**FIGURE S4** Effects of nitrification inhibitors ATU (A), DCD (B), DMPP (C) and NP (D) on the ammonia oxidation activity of *Ca.* Nitrosocosmicus agrestis. Error bars indicate the standard error of the mean for biological triplicates.

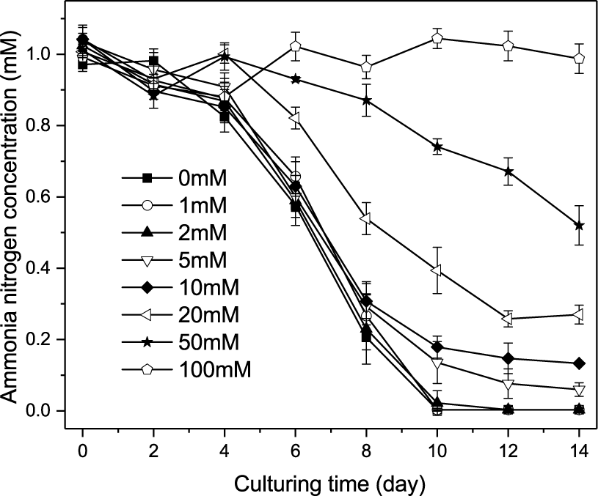

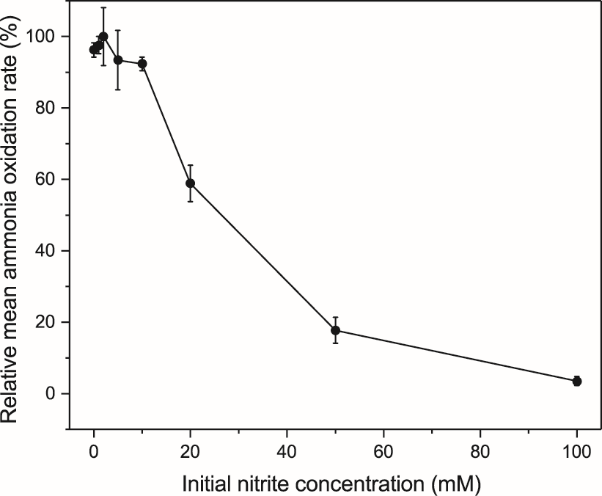

**B**

**A**

**FIGURE S5** (A)Variation of nitrate nitrogen concentration with time at different initial nitrite concentration; (B) the average ammonia oxidation rate of homogenization at different initial nitrite concentration. Error bars indicate the standard error of the mean for biological triplicates.

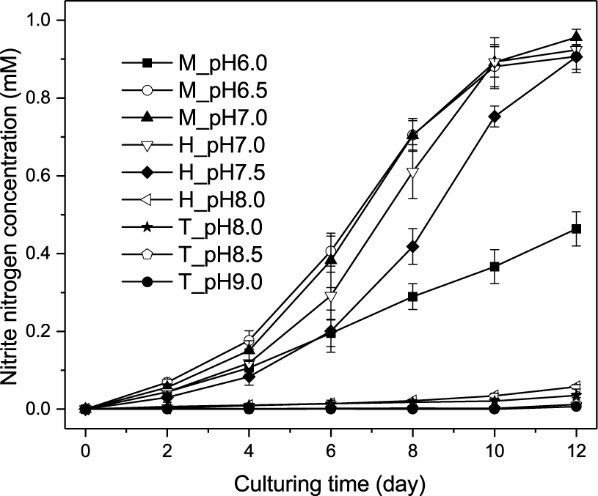

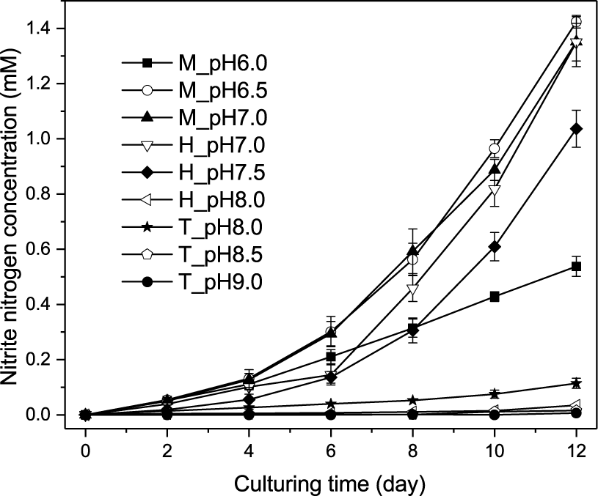

**A**

**B**

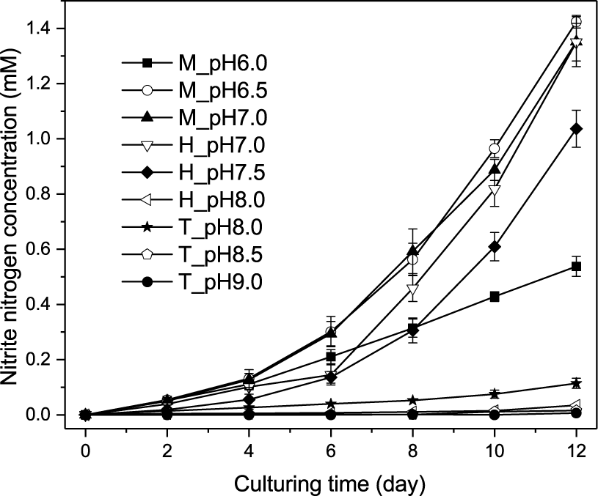

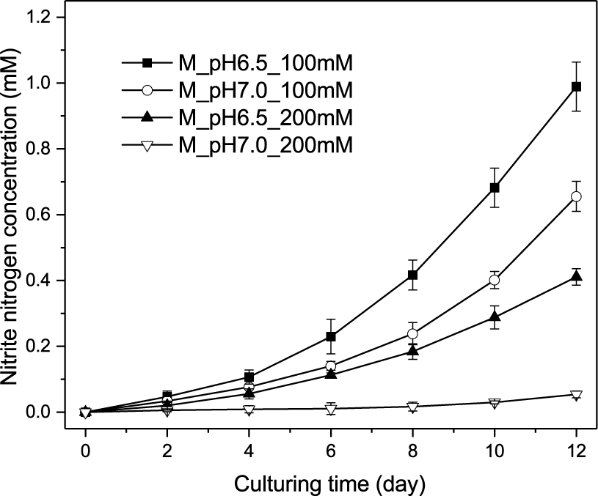

**D**

**C**

**FIGURE S6** Effect of 1mM (A), 10mM (B), 50mM(C), 100mM (D) and 200mM (D) concentration of ammonium on ammonia oxidation activity of strain SS at different pH. Error bars indicate the standard error of the mean for biological triplicates.

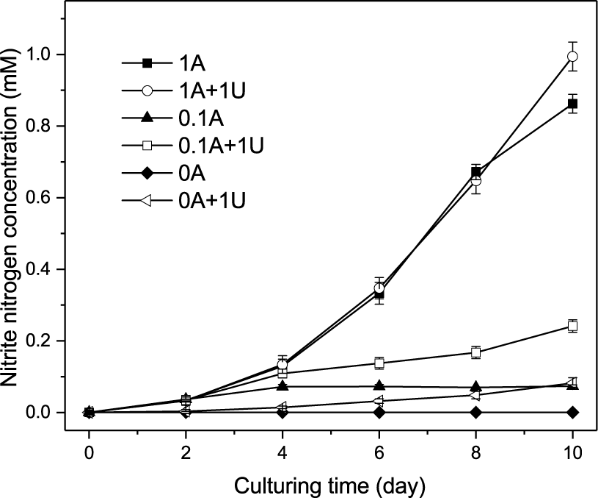

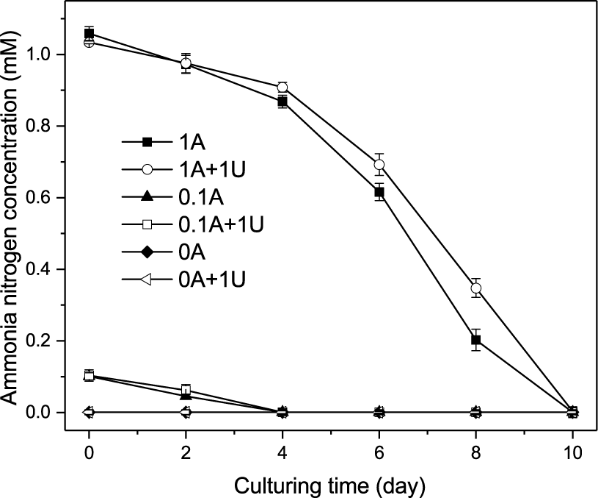

**B**

**A**

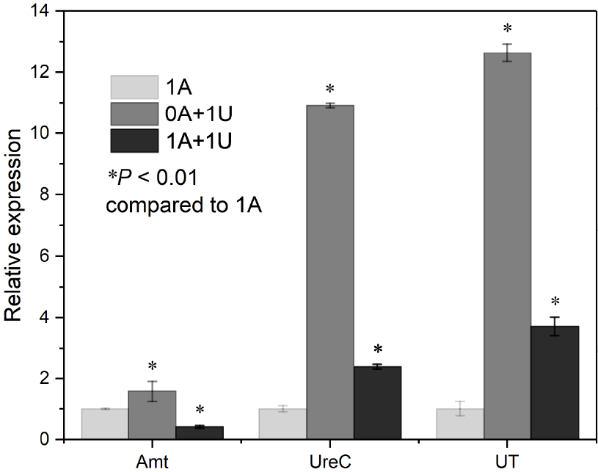

**C**

**FIGURE S7** Effect of urea on nitrite (A) or ammonia (B) concentration in ammonia oxidation process, and relative gene expressions in response to urea (C). 1A, 0.1A, 0A: 1mM, 0.1mM, 0mM NH_4_Cl; 1U: 1mM urea. Amt: Amt ammonium transporter; UreC: urease subunit alpha; UT: urea transporter. Error bars indicate the standard error of the mean for biological triplicates.

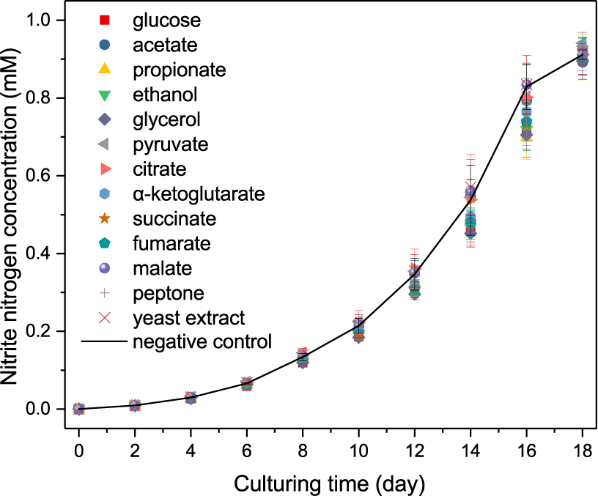

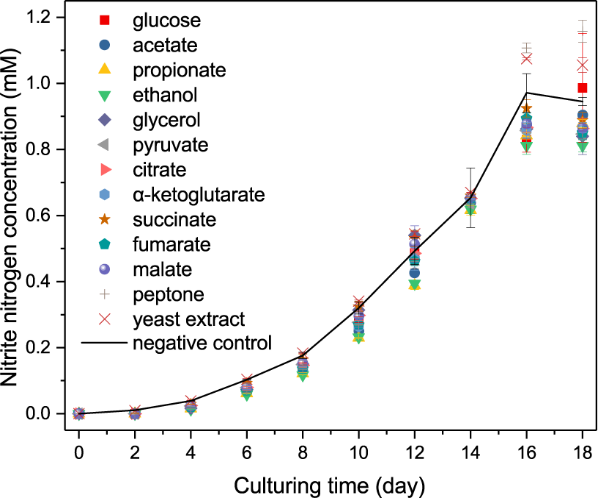

**B**

**A**

**FIGURE S8** Growth of *Ca.* Nitrosocosmicus agrestis amended with organic carbon and incubated with ciprofloxacin- azithromycin (A) or no antibiotics (B). The initial ammonia concentration is 1mM. Error bars indicate the standard error of the mean for biological triplicates.

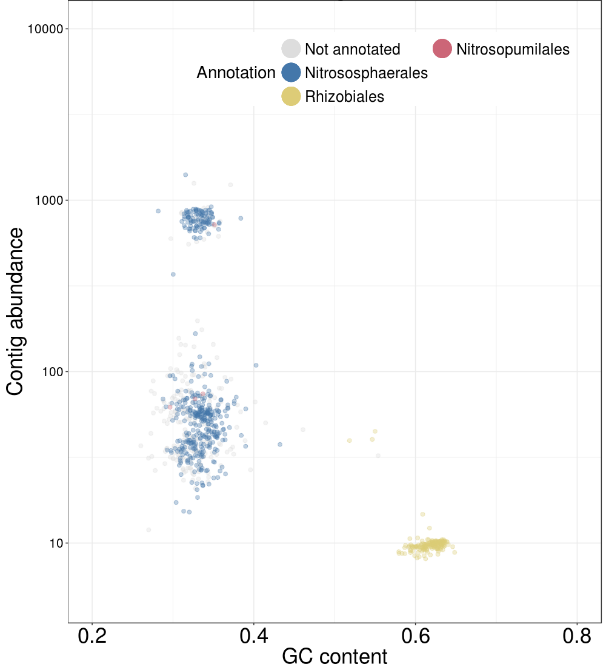

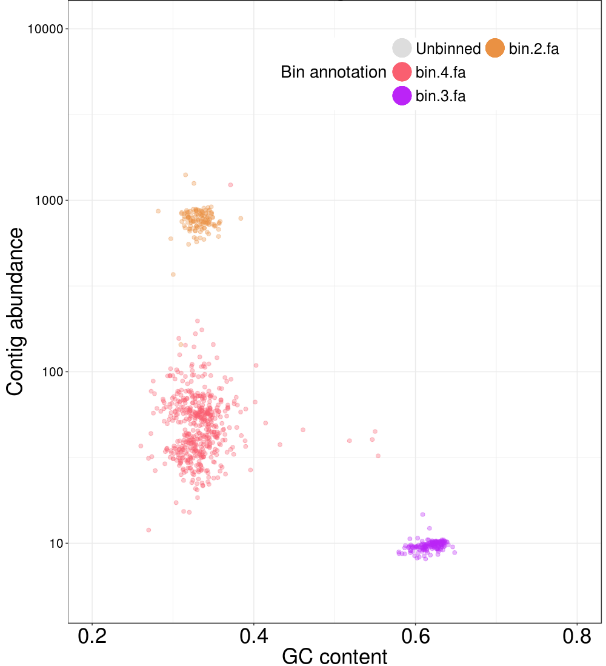

**B**

**A**

**FIGURE S9** Classification and coverage of metagenome contigs (A) and composition of bins from the *Ca*. Nitrosocosmicus agrestis cultures (B); bin2 is the genome of *Ca.* Nitrosocosmicus agrestis.

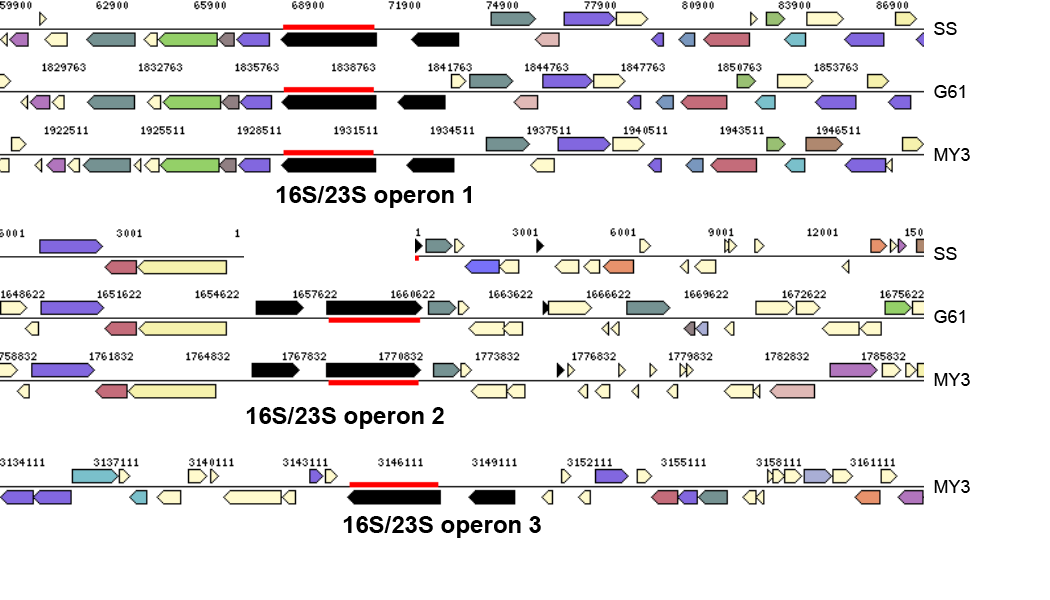

**FIGURE S10** Gene arrangement around 16S/23S operons in the genomes of *Ca.* Nitrosocosmicus. SS: *Ca.* Nitrosocosmicus agrestis SS; G61: *Ca.* Nitrosocosmicus exaquare G61; MY3: *Ca.* Nitrosocosmicus oleophilus MY3.

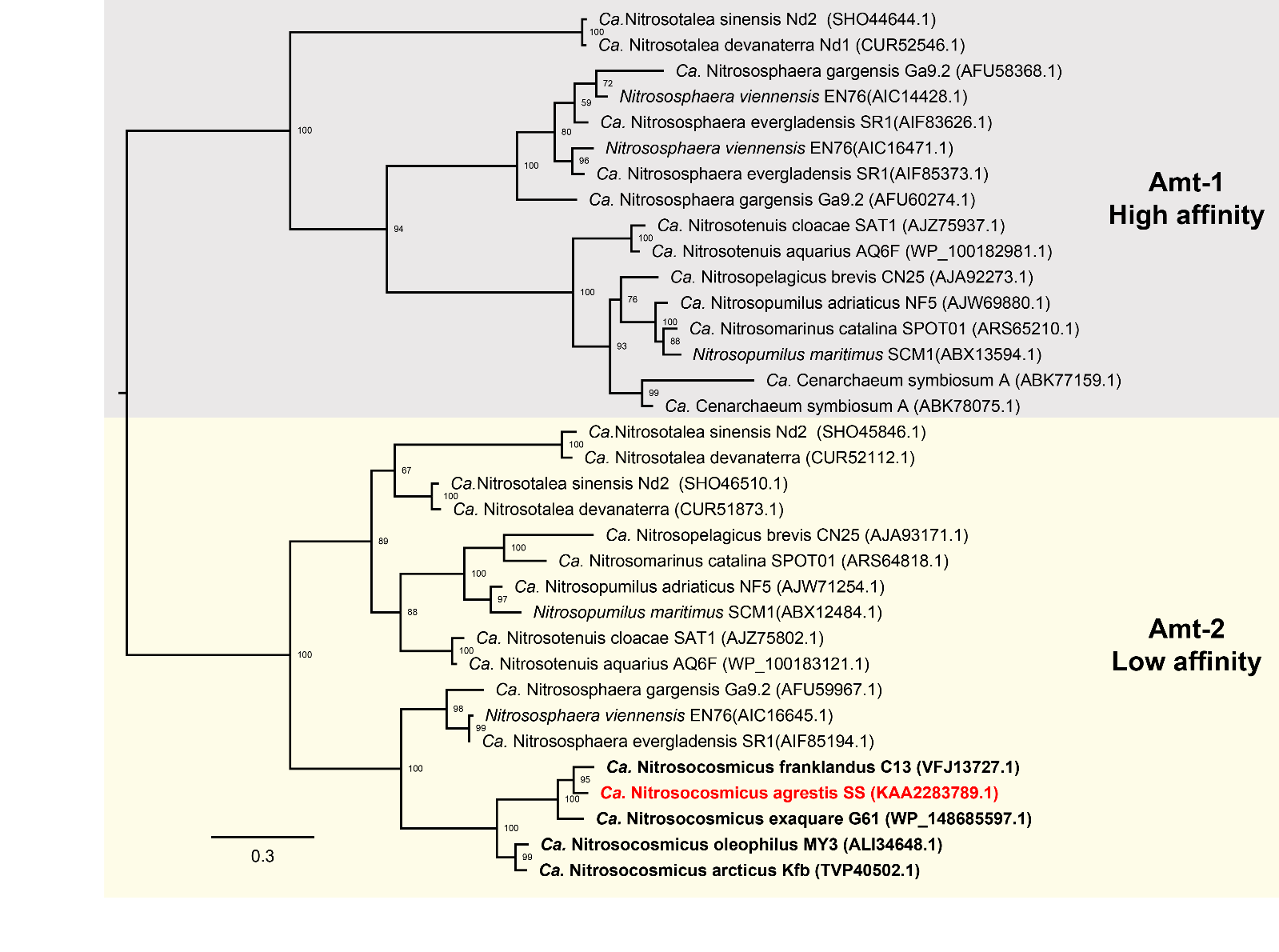

**FIGURE S11** Evolution Analysis of Amt amino acid sequences in the different AOA genomes. Software: iqtree; Method: maximum likelihood; Model: LG+F+R3; bootstrap value: 1000.

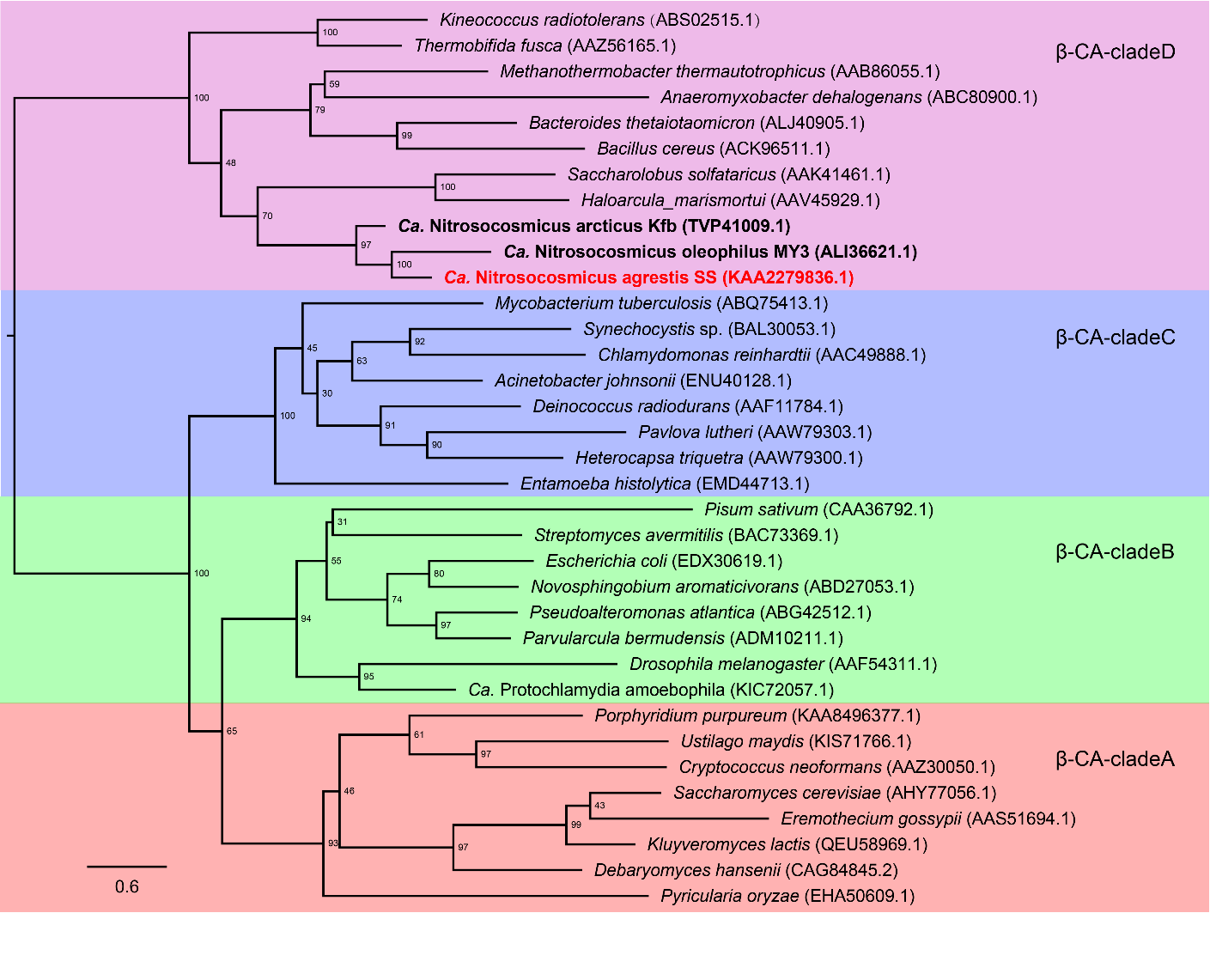

**A**

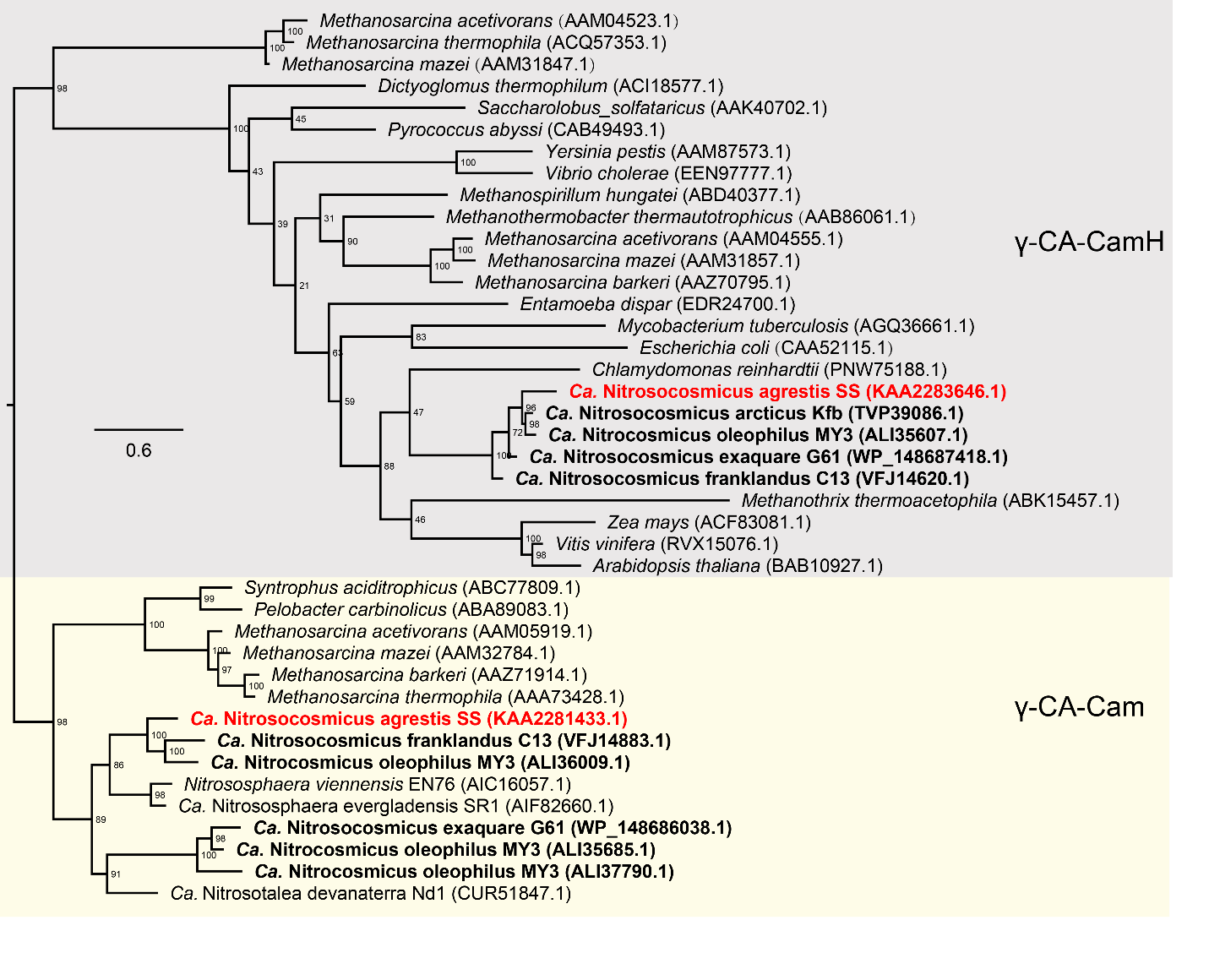

**B**

**FIGURES 12** Evolution analysis of β (A) or γ (B)-carbonic anhydrase based on amino acid sequence. Software: iqtree; Method: maximum likelihood; Model: LG+I+G4; bootstrap value: 1000.

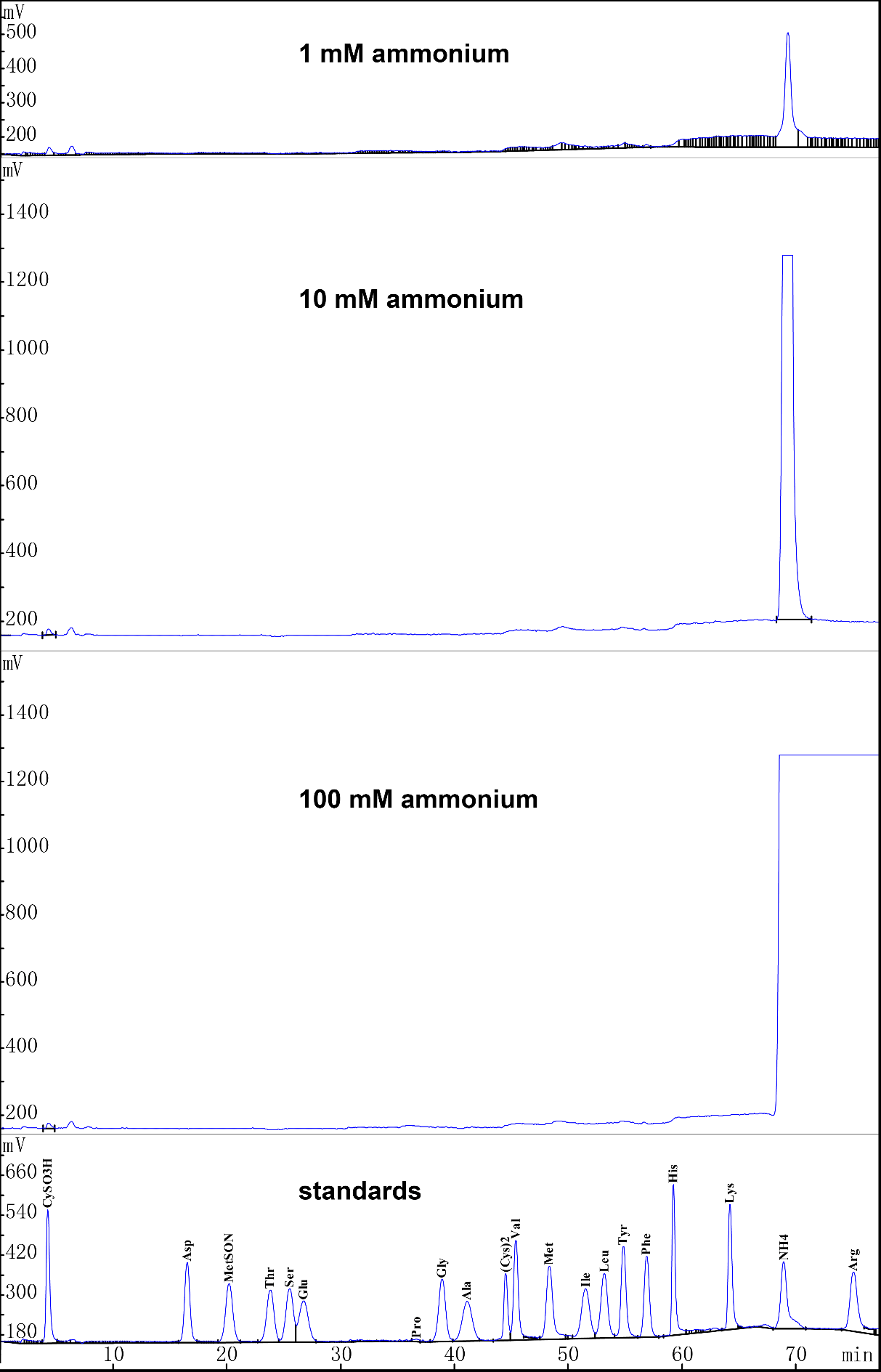

**FIGURE S13** Detection of dissolved free amino acid in the supernatant of the enrichments under different concentrations of ammonium. Except for (Cys)2 of 50 nmol / mL, all other standards are 100 nmol / mL.
